# Phylogenetic signal in primate tooth enamel proteins and its relevance for paleoproteomics

**DOI:** 10.1101/2024.02.28.580462

**Authors:** Ricardo Fong Zazueta, Johanna Krueger, David M. Alba, Xènia Aymerich, Robin M. D. Beck, Enrico Cappellini, Guillermo Carrillo Martín, Omar Cirilli, Nathan Clark, Omar E. Cornejo, Kyle Kai-How Farh, Luis Ferrández-Peral, David Juan, Joanna L. Kelley, Lukas F. K. Kuderna, Jordan Little, Joseph D. Orkin, Ryan S. Paterson, Harvinder Pawar, Tomas Marques-Bonet, Esther Lizano

## Abstract

Ancient tooth enamel, and to some extent dentin and bone, contain characteristic peptides that persist for long periods of time. In particular, peptides from the enamel proteome (enamelome) have been used to reconstruct the phylogenetic relationships of fossil specimens and to estimate divergence times. However, the enamelome is based on only about 10 genes, whose protein products undergo fragmentation *post mortem*. Moreover, some of the enamelome genes are paralogous or may coevolve. This raises the question as to whether the enamelome provides enough information for reliable phylogenetic inference. We address these considerations on a selection of enamel-associated proteins that has been computationally predicted from genomic data from 232 primate species. We created multiple sequence alignments (MSAs) for each protein and estimated the evolutionary rate for each site and examined which sites overlap with the parts of the protein sequences that are typically isolated from fossils. Based on this, we simulated ancient data with different degrees of sequence fragmentation, followed by phylogenetic analysis. We compared these trees to a reference species tree. Up to a degree of fragmentation that is similar to that of fossil samples from 1-2 million years ago, the phylogenetic placements of most nodes at family level are consistent with the reference species tree. We found that the composition of the proteome influences the phylogenetic placement of Tarsiiformes. For the inference of molecular phylogenies based on paleoproteomic data, we recommend characterizing the evolution of the proteomes from the closest extant relatives to maximize the reliability of phylogenetic inference.

## Introduction

### Historical perspective

The survival of endogenous amino acids in fossils was demonstrated in the mid-20th century (Abelson 1954). More recently, access to protein sequence data from long deceased organisms has been achieved with the aid of mass spectrometry methods (Nielsen-Marsh et al. 2002; Cappellini et al. 2014). Since then, the field has grown to propose a set of standards (Hendy et al. 2018; Welker 2018; Hendy 2021; Warinner et al. 2022), and has proven to reliably determine sequence information from samples from as much as 14.8 million years ago (Ma) (Stolarski et al. 2023). The persistence of peptides for millions of years, even from temperate to warm environments, contrasts with the maximum biomolecule age of 2 Ma from ancient DNA (aDNA) under permafrost conditions, which are considered ideal conditions (Kjær et al. 2022).

Even before the development of peptide sequencing, sequence differences in proteins were used for their discriminatory potential in systematics, using e.g. electrophoresis (Johnson and Wicks 1959) or immunological assays (Goodman 1960). Since peptide sequencing preceded DNA sequencing, the first biological sequence comparisons were based on protein sequences. Margoliash (1963), compared cytochrome c sequences of different mammals and baker’s yeast, demonstrating that more closely related taxa share more consensus in homologous amino acid positions than more distantly related groups.

Current advances in sequencing ancient proteins (Cappellini et al. 2019) have created a somewhat similar situation to the first attempts of molecular systematics based on proteins: Due to *post mortem* degradation and - in the case of tooth enamel - low protein abundance in the tissue (Castiblanco et al. 2015), the field of paleoproteomics has focused on the analysis of a very limited amount of peptide sequence data. Again similarly to early days of protein sequencing, scientists have started studying ancient proteomes from a phylogenetic perspective (Welker et al. 2015; Cappellini et al. 2019; Welker et al. 2019; Welker et al. 2020; Madupe et al. 2023).

Given the abundance of tooth remains in the archaeological record, a considerable amount of paleoproteomic research has focused on tooth enamel (e.g. Cappellini et al. 2019; Dickinson et al. 2019; Welker et al. 2019; Froment et al. 2020; Welker et al. 2020; Nogueira et al. 2021; Madupe et al. 2023). Several protein fragments are persistent in mature enamel (Castiblanco et al. 2015). These protein fragments have been successfully used to infer the phylogenetic position of extinct taxa, such as the Pleistocene rhinoceros *Stephanorhinus*, and the extinct hominids *Gigantopithecus blacki*, and *Homo antecessor* (Cappellini et al. 2019; Welker et al. 2019; Welker et al. 2020). However, these studies have also highlighted some of the current challenges of addressing phylogenetic analysis through ancient proteins. The most evident drawback is the limited amount of information due to the short length of the recovered peptides. In particular, the enamel proteome is rather small, comprising less than

15 proteins, which are further enzymatically degraded *in vivo* during enamel formation (Smith et al. 1989), and even more *post mortem*. To date, the combined length of recovered peptides from ancient enamelomes ranges between 456 amino acids (Welker et al. 2019) and 1014 amino acids (Welker et al. 2020). In addition, the peptides from the enamel proteome are not evenly recovered along the protein sequence (Welker et al. 2020), further limiting subsequent analyses.

The main difference to the early days of protein sequencing in the 1960s is that, nowadays, we have much better means to evaluate the phylogenetic signal in a given set of protein sequences. Whole genome sequencing can provide many more informative sites than a whole proteome ever could. This advance has led to a continuous refinement of molecular phylogenies and thus provides a robust reference to compare against protein sequence-based phylogenies. Moreover, protein sequences can be bioinformatically predicted from nucleotide sequences. This enables us to infer protein sequences without the need to sequence the proteins directly.

### Objective

To our knowledge, a comprehensive assessment of the phylogenetic signal present in the enamel proteome has not been made thus far. Here we evaluate the accuracy of phylogenetic reconstructions that can be achieved with fragmentary peptide data, compared to a robust, dated whole-genome phylogeny (Kuderna et al. 2023). We performed several phylogenetic analyses on protein sequences predicted from DNA data that span 16 families of the order Primates (Figure 1). The analysis is based on 14 proteins that have been associated with the enamel proteome (Maas and Dumont 1999; Bartlett et al. 2006; Asaka et al. 2009; Zanolli et al. 2017; Cappellini et al. 2019; Welker et al. 2019; Welker et al. 2020). The proteins analyzed are AHSG, ALB, AMBN, AMTN, AMELX, ENAM, MMP20, ODAM, SERPINC1, TUFT1, COL1A1, COL1A2, COL17A1, and COL2A1. The selection of these proteins was mainly driven by the availability of experimental proteomic and genomic data. Other proteins associated with tooth enamel, such as KLK4, may play a key role in enamel formation (Yamakoshi 2006), but barely leave behind any peptides that can be experimentally recovered in paleoproteomic studies (Cappellini et al. 2019; Welker et al. 2019; Welker et al. 2020; Madupe et al. 2023). Similarly, AMELY is considered enamel-specific, but since it is encoded on the difficult-to-sequence Y chromosome, there is little genomic reference data available. COL1A1, COL1A2, and COL2A1 have been included in this study, although they are not canonically considered to be part of tooth enamel, because they are occasionally co-extracted from dentin fragments still attached to ancient enamel samples processed for paleoproteomic analysis (Madupe et al. 2023), or because they are recovered in experiments targeting bone or dentin on younger fossils (Welker et al. 2015; Chen et al. 2019; Presslee et al. 2019). Lastly, these collagens are of great importance for the peptide mass fingerprinting method conventionally called “zooarchaeology by mass spectrometry”, short “ZooMS” (Buckley et al. 2009; Naihui et al. 2021).

**Figure 1:**
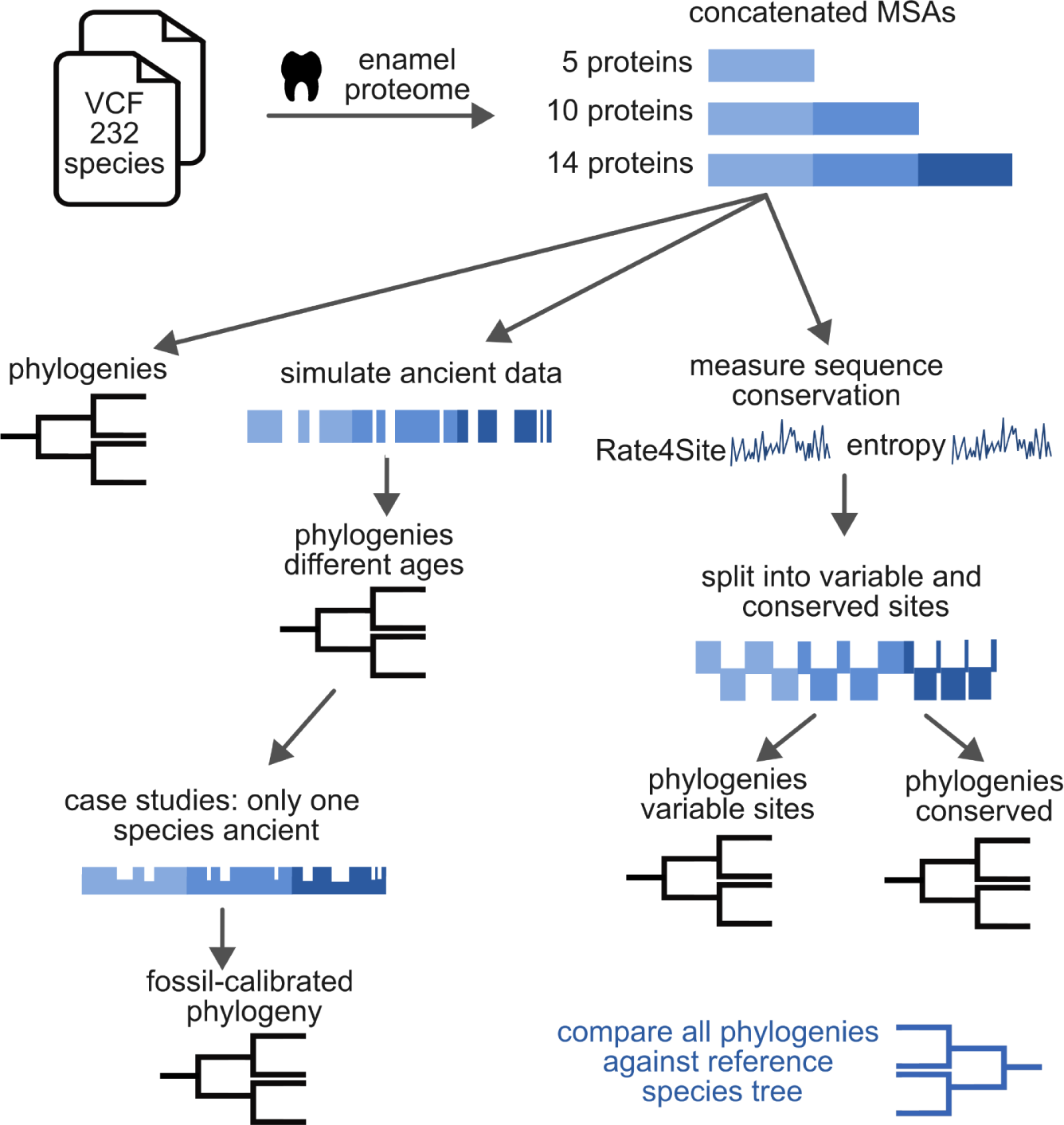
Overview of workflow. All primate genomic data stems from previously published VCF files (Meyer et al. 2012; Prado-Martinez et al. 2013; Prüfer et al. 2014; Xue et al. 2015; De Manuel et al. 2016; Mallick et al. 2016; Nater et al. 2017; Prüfer et al. 2017; Kuderna et al. 2023). From these files we predicted the sequences of 14 tooth enamel proteins. The sequences of these proteins were aligned and concatenated into larger MSAs, combining different proteins. One version contains the 5 proteins that have been experimentally verified in several studies (Cappellini et al. 2019; Welker et al. 2019; Welker et al. 2020; Madupe et al. 2023), the 10 protein version contains 5 additional proteins that may be found in enamel, and the 14 protein version contains 4 additional collagen sequences. We performed phylogenetic analysis on the full sequence of these concatenated MSAs (excluding signal peptide). We also simulated ancient, fragmentary data for different degrees of fragmentation by eliminating sites in the MSA equivalent to data loss seen in ancient samples. Subsequently, we performed phylogenetic analysis on these MSAs, either with all or only one species fragmented. Moreover, we quantified the degree of variability of amino acid sites across primates using Shannon entropy and Rate4Site (Pupko et al. 2002). This way, sites of the MSAs could be categorized into “conserved” or “variable” and phylogenetic analysis could be performed on each of those sets of sites. All phylogenetic trees resulting from the analyses of this project were compared to a genomic data-based reference tree (Kuderna et al. 2023).

Phylogenetic analysis was performed on the full-length translated sequences of the 14 proteins listed above. A further analysis was carried out only with peptides corresponding to the protein regions typically captured in paleoproteomic studies with the aim of simulating the limited amount of data in paleoproteomic studies. We further searched for segments in the protein sequences that are most phylogenetically informative. The results of these analyses will inform future paleoproteomic studies by indicating which peptides should have priority in experimental recovery, but also by setting realistic expectations for the discriminatory power of these sequences in subsequent phylogenetic studies. Lastly, we discuss the implications of these findings and related factors, such as possible dependencies between the studied loci, when using ancient enamel peptides for evolutionary studies.

## Materials and Methods

### Dataset

The primate DNA sequences stem from 718 Variant Calling Format files (VCFs) from whole genome sequence data which were analyzed along with publicly available DNA sequences of the outgroup taxon *Tupaia belangeri chinensis*. In total, this represented 719 individuals: 135 great apes (Prado-Martinez et al. 2013; Xue et al. 2015; De Manuel et al. 2016; Nater et al. 2017) mapped to hg19; 561 individuals spanning 16 primate families (including more great apes) mapped against 31 primate genomes as listed in the supplementary information section 1 (Kuderna et al. 2023); 19 modern humans from the Simons Genome Diversity Project (Mallick et al. 2016) and three extinct hominins (Meyer et al. 2012; Prüfer et al. 2014; Prüfer et al. 2017), all which were mapped to hg19, and publicly available protein sequences of *Tupaia* as the outgroup (Fan et al. 2013). Sequences of *Nomascus leucogenys* and *Pongo tapanuliensis* were later excluded due to low quality. Sequences of Neanderthal and Denisovan were only included in one case study (“Neanderthal case”).

### Amino acid sequence translation and multiple sequence alignments

For all 14 genes under study, we restricted our analyses to the canonical isoforms from the human hg38 annotation (Ensembl) (supplementary table S1) to ensure comparable sequences across species. In addition to their own annotation, the hg38 annotation was projected onto the 31 reference genomes of the 561 primates from Kuderna et al. (2023). Using the liftOver tool (Hinrichs et al. 2006) with default parameters (supplement section 1), we obtained GTF-files of the projected coding sequence (CDS) coordinates for each of the 31 reference genomes. For the 157 representatives of the family Hominidae (i.e., great apes, and fossil and modern humans) that were originally mapped to hg19, the hg19 human annotation was used for simplicity, because the annotations of the genes of interest did not show any significant differences between the hg19 and hg38 annotations. The VCFs were used to integrate genomic variants in the CDS of interest using samtools (Li et al. 2009) and bcftools (Li 2011) (supplement section 1). For each individual, the resulting CDS were translated to proteins through in-house python scripts based on the standard genetic code. Low-quality regions at the DNA level were represented as “N”, and affected codons masked as an “X” in the amino acid sequence.

The resulting translations were grouped by protein and aligned with MAFFT v7.520 (Katoh et al. 2002). Alignments were trimmed using trimal 1.2rev59 (Capella-Gutiérrez et al. 2009) (for parameters see supplement section 1). The resulting alignment files per protein were manually explored to identify sequence segments that differed markedly from the variation observed in the alignments. These segments were characterized as spurious variation, and were caused by frameshifts due to indels which were shared by individuals mapped to the same reference genome. In order to recover the frame that removed or minimized this spurious variation, the coordinates of this variation were registered and used to edit the DNA data (in-house Python code) and retranslated. The newly translated protein sequences were realigned and trimmed, masking any remaining probable spurious variation residues with a “?”.

Besides LiftOver-based annotations, an original annotation existed for each of the 31 reference genomes of the above mentioned 561 species (Kuderna et al. 2023). About half of these reference genomes have annotations that were previously published and have been achieved in various ways (see accessions in Kuderna et al. 2023). The other half stems from Shao et al. (2023), and has been annotated with a combination of de-novo and homology-based strategies. In some cases, the predicted protein sequence from the original annotation resulted in a higher quality protein model than LiftOver-based annotations (less premature truncation, less spurious variation), but in other cases the opposite was true. To determine the best annotation for each protein in each reference genome, we aligned all predicted protein sequences (original and LiftOver-based annotations) against the human canonical protein and manually inspected the alignments. The protein model that yielded the fewest number of gaps was kept for further analysis.

Different sets of MSAs were concatenated, comprising groups of 5, 10, and 14 proteins. The 5 protein concatenation consists of AMBN, AMELX, AMTN, ENAM and MMP20, proteins that are an integral part of the enamel structure and have been consistently identified from fossil teeth in previous studies (Cappellini et al. 2019; Welker et al. 2019; Welker et al. 2020; Madupe et al. 2023). The 10 protein concatenation represents a larger, non-collagenous enamel proteome by adding AHSG, ALB, ODAM, SERPINC1, and TUFT1. The 14 protein concatenation also included 4 collagens: COL17A1, COL1A1, COL1A2, and COL2A1. For subsequent phylogenetic analyses, the signal peptide sequence was removed from every protein sequence, given that it is never recovered in paleoproteomic experiments. If not otherwise stated, in the following, “MSA’’ always refers to a concatenation of different sets of proteins of interest. Variable and parsimony-informative sites were assessed using MEGA11 (Molecular Evolutionary Genetics Analysis v. 11) (Tamura et al. 2021).

### Shannon entropy and Rate4Site calculations

Shannon entropy is a measure that can be applied to MSAs to quantify the degree of variability at each given homologous site. It is agnostic to physico-chemical similarities and substitution rates between amino acids. It was calculated with a moving average of 20 (https://gist.github.com/jrjhealey/Shannon.py) on the concatenated MSA of 14 proteins with one individual per species. Rate4Site (Pupko et al. 2002) is a tool used to calculate conservation scores in homologous amino acid sites. For the same MSA, Rate4Site scores were calculated using default options and setting the concatenated *Tupaia belangeri chinensis* proteins (outgroup) as reference sequence. Gaps in *Tupaia* proteins were first filled in with the consensus sequence, since the Rate4Site tool will omit sites with an incomplete reference. As for the calculation of the Shannon entropy, a moving average of 20 was used to calculate all Rate4Site scores. Alternatively, for the estimation of evolutionary rates in these proteins across mammals, Rate4Site scores were calculated on a concatenated MSA of 22 species (for list of species and sequence IDs see supplementary table S4). The protein sequence data was downloaded from UniParc using the ProteoParc v1.0 tool (https://github.com/guillecarrillo/proteoparc), which facilitates download of proteomes from a given clade and curation of fasta files. We selected a set of species that had a mostly complete sequence for each gene and that represented most clades across the group of mammals. Rate4Site scores were calculated using default parameters, setting the reference sequence to *Homo sapiens*.

### Examination of the effect of intraspecies variation

The concatenated MSA of all 719 individuals was randomly downsampled to contain exactly one individual per species (1,000 iterations) using in-house scripts. Hence, each of these MSAs comprises the concatenated proteins from 232 primate individuals and one individual of the outgroup *Tupaia belangeri chinensis*. To measure the effect of different combinations of individuals on the downstream analyses, from each MSA, we calculated the Shannon entropy, Rate4Site scores, and a maximum likelihood (ML) phylogenetic tree using IQ-TREE v 1.6.12 (Nguyen et al. 2015). The Shannon entropy and Rate4Site scores were compared to those of the most complete MSA (i.e. with least masked positions) by subtracting the values at each respective amino acid position and calculating the mean of the absolute difference for each downsampled MSA. To estimate the magnitude of these mean differences, they were transformed into a percentage of the possible ranges of values in the reference MSA (i.e. divided by the range between min. and max. entropy or R4S score). The accuracy of the topology of the 1,000 ML trees was assessed by measuring the Robinson– Foulds distance (“RF-distance”, topology only) to the reference tree (Kuderna et al. 2023) in R v.4.3.1 using the package ‘phangorn’ v.2.11.1 (Schliep 2011).

### Phylogenetic analysis

For any further downstream analysis, the MSA of all 719 individuals was downsampled to one individual per species (with the most complete sequence), yielding a total of 233 individuals. All phylogenetic analyses were performed using ML with IQ-TREE v1.6.12 (Nguyen et al. 2015) including the Shimodaira Hasegawa approximate likelihood-ratio test (SH-alrt), for 5000 iterations with ultrafast bootstrap approximation. The evolutionary model of each of the individual proteins was obtained through ModelFinder (Kalyaanamoorthy et al. 2017). For a detailed description of the code and parameters see section 2 in the supplementary information. In addition, for the complete protein sequence (except the signal peptide) of all three concatenations (5, 10, and 14 proteins), phylogenetic trees were also calculated using Bayesian analysis performed using MrBayes v.3.2.7a (Ronquist and Huelsenbeck 2003). Each Bayesian analysis was run for 3 million generations, with a burn-in of 25%. For all trees, the distance to the reference species tree (Kuderna et al. 2023) was assessed via RF-distance using the R package ‘phangorn’. Next, we calculated phylogenetic trees using the above-mentioned parameters for different subsets of amino acid positions. The rationale for building these subsets is described in the following sections.

### Ancient sequence reconstruction of enamel peptides from fossil specimens

Ancient peptide sequences were isolated from the tooth enamel of fossil equids and deinotheriid proboscideans of different ages. The former include specimens of *Equus* cf. *ferus* (IPS87498, 138 mg enamel powder, and IPS87522, 820 mg) from the Late Pleistocene of La Carihuela (probably <100 ka), *Equus* cf. *altidens* (IPS137786, 169 mg) from the Early Pleistocene of Vallparadís layer EVT7 (0.9-0.8 Ma) (Aurell-Garrido et al. 2010; Madurell-Malapeira et al. 2010), and *Hippotherium* cf. *primigenium* (IPS98842, 50 mg) from the Late Miocene of Can Llobateres 1 (9.8 Ma) (Casanovas-Vilar et al. 2016; Arranz et al. 2023). Considering that a detailed study of these fossil samples is needed, at the present time we use the open nomenclature for the specimens used. In turn, deinotheriid specimens correspond to *Deinotherium giganteum* (IPS28029, 80 mg) from Can Llobateres (see above) and *Deinotherium levius* (IPS121827, 130 mg) from the Middle/Late Miocene of Abocador de Can Mata locality ACM/C8-A3 (11.6 Ma) (Alba et al. 2022). All the fossil specimens are housed in the Institut Català de Paleontologia Miquel Crusafont, Sabadell, Spain.

Enamel samples were precisely extracted using a rotary tool with a diamond disc and a slow-speed drill (Dremel®); traces of dentin adhered to the enamel were removed with a scalpel and fiberglass pencil. Ancient peptide sequences were isolated from the enamel pieces in a dedicated clean room following published protocols (Cappellini et al. 2019; Welker et al. 2019), using trifluoroacetic acid (TFA) as the demineralizing agent. The solubilized peptides were immobilized on a C18 membrane STAGE tip (Rappsilber et al. 2007) and washed with 5% v/v formic acid. Elution followed with a 5% v/v formic acid 50% v/v acetonitrile solution. The eluted peptides were subjected to reverse phase nano liquid chromatography coupled with tandem mass spectrometry. Samples were analyzed using an Orbitrap Eclipse mass spectrometer (Thermo Fisher Scientific, San Jose, USA) coupled to an EASY-nLC 1200 (Thermo Fisher Scientific, San Jose, USA). More details on the run on this instrument are described in supplementary information section 1.4. As negative controls, extraction blanks were processed together with the ancient samples during peptide extraction. In addition, injection blanks were injected into the mass spectrometer, between the single injections of the samples and extraction blanks.

The ancient peptides were identified in iterative reference database searches using MaxQuant and MaxNovo. The databases were built from public repositories using the ProteoParc v1.0 tool. A list of the proteins in the databases and database search parameters can be found in supplementary information section 1.5 (supplementary table S2). The resulting ancient reconstructed sequences were used to inform the creation of subsets of the MSA.

### Reducing alignments to simulate ancient peptide sequences

To date, there is only a limited amount of information on how enamel proteins degrade *post mortem* over large time scales. Nevertheless, we created a simple model of peptide fragment degradation by inspecting most publicly available enamel proteomes, combined with new data from this study. Ancient protein sequences, also known as ancient sequence reconstructions, ASRs, (Cappellini et al. 2018), from tooth enamel of various mammals were downloaded from publicly available data (Welker et al. 2015; Cappellini et al. 2019; Chen et al. 2019; Presslee et al. 2019; Welker et al. 2019; Welker et al. 2020) or sequenced at the Institute for Evolutionary Biology and the Centre for Genomic Regulation (CRG) in Barcelona (see section above). The ancient sequences were aligned using MAFFT v7.520 to the corresponding human reference protein from UniProt (supplementary file 3). The alignments were manually curated because the highly fragmentary nature of the sequenced ancient peptides can cause misalignment at non-homologous positions. These curated alignments were added to the predicted protein sequences of this study, using MAFFT v7.520 with the -- add and --keeplength option, in order to keep the number of columns in the predicted sequence MSA the same. Different stages of *post mortem* protein sequence fragmentation were modeled by removing sites (i.e. columns) in the MSA of each protein. The older the modeled fragmentation stage, the more sites were removed in a non-randomized manner. For the model we assumed heterogenous *post mortem* survival times across all sites of each protein for two reasons: First, because the enamelome is already enzymatically cleaved *in vivo*, and second because different peptides may have different physico-chemical properties that influence their chemical breakdown. To understand the approximate patterns of this heterogeneity, we assessed in the MSAs how often each site in a protein could be experimentally recovered. To date, the amount of available ancient protein sequences is not sufficient to use statistical methods to model the process of fragmentation across millions of years. Instead, we followed the rationale of reducing the amount of sequence information similar to what we observed in sequences of a certain age range. These ages give rise to the eponymous fragmentation stages. For instance, the MSA of the fragmentation stage “100 ka” has 3884 sites, which are reduced to 1139 sites in the fragmentation stage “1-2 Ma”. The more coverage a single site has, the longer is its anticipated survival. Reducing the MSA to those positions was done using an in-house python script.

A list of all positional information, in relation to the single gene MSAs before concatenation, can be found in supplementary file 4. Note that the sample ages that describe the different fragmentation stages (“100 ka”, “1-2 Ma”, “5 Ma”, “10 Ma”) are based on the actual age of each sample and that most of them (in case there was metadata about the exact location) stem from sites with annual average temperatures higher than 10 °C. The fragmentation stage in samples of similar age might be different depending on its environment. For the stage “100 ka”, a rather large coverage of collagens is anticipated because, at this fragmentation stage, additional sampling of dentin or bone may be possible. For the stage “5 Ma”, experimental data is not available, so that this stage is an intermediate between “1-2 Ma” and “10 Ma”. We could not find any public peptides that stem from TUFT1, nor could we confidently sequence them. Phylogenetic analysis with ML was performed on the four subset MSAs, and the resulting topologies were compared against the reference tree with RF-distance.

### Case studies of simulated ancient samples

With the aim of simulating typical phylogenetic inference with paleoproteomic data, two phylogenetic analyses were performed as two case studies (“Neanderthal case” and “Chimpanzee case”). In these two scenarios, the “100 ka” fragmentation pattern was used for the “Neanderthal case”, and the “5 Ma” pattern for the “Chimpanzee case”, aiming to mimic the fragmentary pattern that could be recovered from actual fossils of both cases after their split from their most recent extant sister group. The objective of these two analyses was to observe if the fragmented data still allowed the individuals to be positioned correctly in the phylogenetic tree. The reference data for the “Neanderthal case” consisted of a concatenation of the set of 14 protein (full-lengths) of five individuals from each of the hominid species, including five *Homo sapiens* individuals, and one *Hylobates lar* individual as an outgroup. The three Neanderthal/Denisovan sequences with the “100 ka” fragmentation pattern were added to this scaffold. The “Chimpanzee case” also comprised 14 concatenated proteins of five individuals per hominid species, including five *Homo sapiens*, but excluding *Pan troglodytes*, and using *Hylobates lar* again as an outgroup. Three simulated *Pan troglodytes* sequences with the fragmentation pattern of “5 Ma” were then added to this scaffold. ML phylogenetic analysis was performed on both of these case studies. For more details see the supplementary information section 5.

### Estimation of divergence times

To compare the effect of limited proteomic data on the performance of divergence time estimation, one of the above described case studies underwent further analysis in this regard. The estimation of divergence times was methodologically as similar as possible to the analysis used to generate the reference tree. Consequently, MCMCtree from the PAML program package v. 4.10.6 (Yang 2007) was used for fossil-calibration-based analysis. To format the required protein sequence input, the MSA of the “Chimpanzee case” was pruned to one individual per species, keeping the individuals with the most complete sequences. The same pruning was applied to the reference tree. All fossil calibrations are from de Vries and Beck (2023), and the prior distributions were taken from Kuderna et al (2023). In the control file of MCMCtree, all parameters were set following “tutorial 4” in the “MCMCtree tutorials’’ (dos Reis and Yang 2017) with some modifications to perform an analysis as similar as possible to that of Kuderna et al (2023): The time scale was set to 1 million years. For the approximate likelihood calculation (usedata=2) under the “correlated rates’’ clock model (clock=3), we ran 10 replicates of 1 million generations, sampling every 50 generations. 10% were discarded as burn-in. Convergence of runs was confirmed in TRACER v. 1.7.2 (Rambaut et al. 2018). The raw Markov Chain Monte Carlo (MCMC) results (mcmc.txt) were further analyzed in R using the packages ‘coda’ v.0.19.4 (Plummer et al. 2006) and ‘reshape’ v.0.8.9 (Wickham 2007) for calculating a combined 95% highest posterior density (95% HPD). The package ‘RcolorBrewer’ v.1.1.3 (Harrower and Brewer 2003), and the package ‘ggplot2’ v.3.4.3 (Wickham 2016) were used for plotting. Effective priors were calculated in a separate run (usedata=0).

### Reducing alignments to highly conserved or variable sets

For subsequent analysis the MSAs were separated into sections of higher or lower conservation. We used two methods (Rate4Site and Shannon entropy) to measure variability of each site in the MSA. The Rate4Site score and Shannon entropy values were calculated for each protein and normalized to a mean of one. The MSAs of each protein were then subset by values equal or higher than one and lower than one and concatenated into a long MSA, as in all other analyses. For the two metrics, this resulted in four different types of concatenated MSAs “Shannon variable”, “Shannon conserved”, “Rate4Site variable”, “Rate4Site conserved”. This applied to all three concatenations (5, 10, and 14 proteins) resulted in a total of 12 MSAs. Phylogenetic analysis with ML was performed on each of them. The resulting tree topologies were compared to the trees resulting from the full-length proteins and to the reference tree using RF-distance.

### Evolutionary Rate Covariation scores

The degree of evolutionary co-variation of the set of 14 genes was estimated using Evolutionary Rate Covariation (ERC) analysis (Clark et al. 2012). The ERC for 19,137 orthologous genes from 120 mammalian species was calculated using the R code available at https://github.com/nclark-lab/erc. The covariation in relative evolutionary rates for each gene pair was calculated using only the branches that are shared between the two genes. The raw correlations were then Fisher transformed, normalizing for the number of branches that contributed to the correlation. In R v.4.3.1, significance was estimated via permutation analysis using the mean as test statistic and 10,000 permutations. The results were plotted in R using the package ‘ggplot2’.

## Results

### Shannon entropy and Rate4Site

Both Shannon entropy and Rate4Site can be applied to measure the degree of protein sequence conservation. While Rate4Site accounts for the different likelihoods of substitution during sequence evolution, Shannon entropy values are agnostic to any evolutionary or physico-chemical constraints.

The degree of sequence conservation and evolutionary rates were examined on a set of 14 proteins from 232 primate species and one non-primate outgroup (*Tupaia*). The analysis was performed on a concatenation of MSAs of each protein into one large multi-protein MSA. Each species was represented by the individual that had the most complete sequence data (i.e. fewest gaps or masked positions). As sequence data of more than one individual was available for many species, we estimated the impact of intraspecies variation by calculating Shannon entropy, Rate4Site scores, and phylogenetic trees from the 14-protein-MSAs created by randomly picking one individual per species (n=1,000 iterations) (supplementary figure S1). All trees were compared to a reference species tree that is based on high quality genomic data (Kuderna et al. 2023) by measuring the Robinson-Foulds distances (RF-distances) between them. All RF-distances to the reference tree range between 82-156, showing median (110) and mean (112.7) values that are very close to the RF-distance between the phylogeny based on a single representative individual per species (the one with most complete sequence) and the reference tree (108). The mean differences of Shannon entropy at all given sites range from 0.77% to 2.81% of the possible range of values, with a mean difference of 1.71% (mean difference in entropy value 0.05). The mean differences of Rate4Site scores at all given sites range from 0.50% to 1.36% of the possible range of values, with a mean difference of 0.92% (mean difference in Rate4Site score 0.04). From these results, we assumed that the concatenated MSA of one individual per species with the most complete sequence is representative and continued all downstream analyses with this set of individuals (supplementary file 3).

Shannon entropy values and Rate4Site scores both demonstrate that sequence diversity and evolutionary rates vary across the length of each protein sequence (Figure 2). In particular, collagens (except some sites in COL17A1) evolve at a slower rate than non-collagen proteins (Figure 2B). Rate4Site scores roughly correlate with Shannon entropy values (Figure 2, Pearson correlation ρ = 0.54, p-value < 2.2e-16). About 4% of all sites are residues with particularly high evolutionary rates (r4s score > 2) that also fall into the regions that could be experimentally recovered in ancient samples (Figure 3A). This is the case in particular in ALB, AMELX, AMBN, and ENAM. In most other proteins, such as AHSG, AMTN, COL17A, MMP20, ODAM or TUFT1, the regions of particularly high evolutionary rates correspond to peptides that have not yet been experimentally recovered. On the other extreme, some regions of high sequence conservation levels stand out. In ENAM, there is a stretch of 49 highly conserved amino acids (corresponding to the positions 191-239 in UniProt ID Q9NRM1) that carries two phosphorylation sites. The region falls into the 32 kilodalton (kDa) cleavage product of ENAM (Ozdemir et al. 2005), which belongs to peptides that could be experimentally recovered in deep time (Figure 3) (Welker et al. 2020). MMP20 also displays larger regions of highly conserved amino acids that fall into the experimentally recovered sequences. One of those regions (corresponding to UniProt ID O60882, positions 174-254) lies around the active center (position 227) and its surrounding inorganic ion binding regions; another one lies around positions 330-483, a region whose ends are connected via a disulfide-bridge. Other longer experimentally recovered regions of relatively highly conserved sequences belong to AMBN, COL1A1, COL1A2, COL17A1, and SERPINC1.

**Figure 2:**
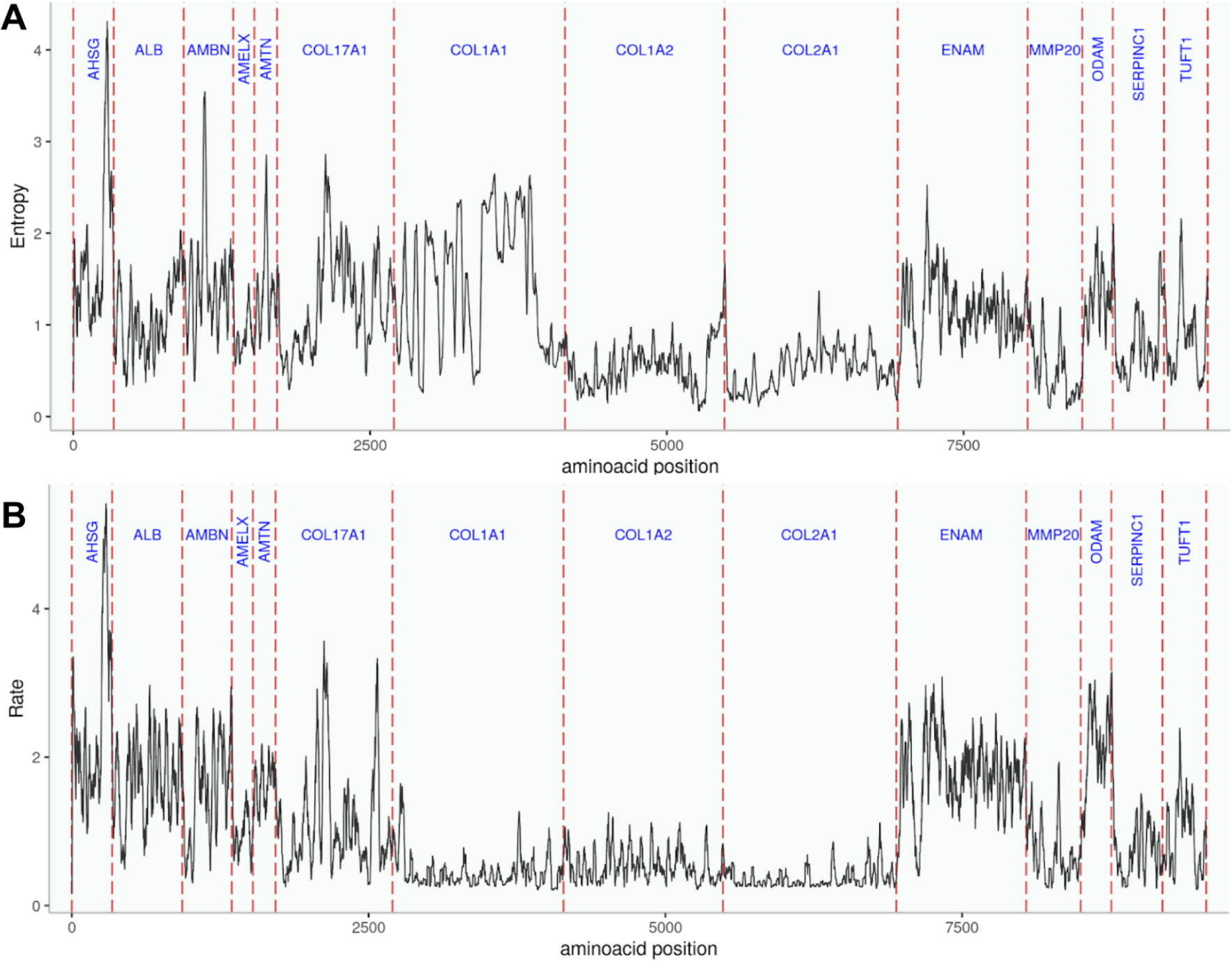
Evolutionary rates and sequence diversity estimated by (A) Shannon entropy and (B) Rate4Site scores for a concatenation of all 14 proteins. Collagens evolve at a slower rate than all non-collagen enamel proteins. COL17A1, which is the only collagen known to be an essential part of tooth enamel (Asaka et al. 2009), has an evolutionary rate and degree of conservation more similar to the non-collagen enamel proteins. COL1A1 displays elevated Shannon entropy values because many sequences have masked or missing positions that the software interprets as diversity.

**Figure 3:**
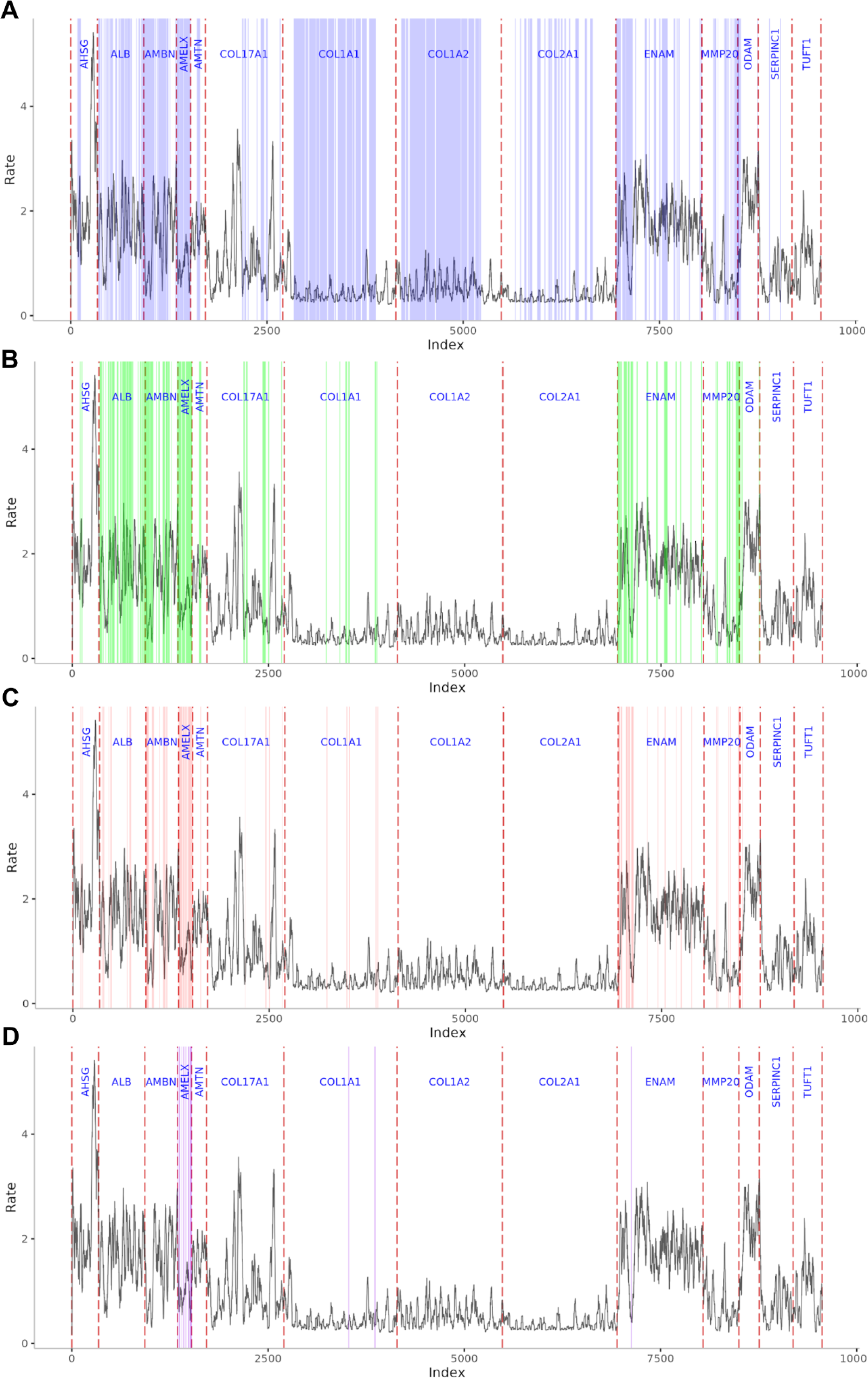
Evolutionary rates in the context of potential ancient sequence coverage across different time scales. The colorful shaded areas represent areas that have experimental support for being able to be retrieved from fossil tooth enamel samples. The names of the stages represent the ages of the samples they are based on, which all stem from moderate to tropical climate zones. They may not necessarily reflect the stage of degradation of any sample at this given age. A - “100ka” (collagens may be retrieved from dentin or bone); B - “1-2 Ma”; C - “5 Ma” (no direct fossil evidence, extrapolated between “1-2 Ma” and “10 Ma”), D - “10 Ma”

To examine if the above described patterns also hold beyond primates, Rate4Site scores were calculated on a set of 22 species from different taxonomic groups across mammals (supplementary figure S4, for list of species see supplementary table S4). The general pattern of slower evolutionary rates in collagens is consistent across mammals, in particular for COL1A1, COL1A2, and COL2A1. The area of the 32 kDa fragment in ENAM is not as strongly conserved as within primates, whereas the region around the active center in MMP20 shows a persistently low evolutionary rate. Across mammals, the N-terminus of AMELX displays a higher degree of conservation as within primates. In contrast, the C-terminus of AMTN, and the N-terminus of COL1A1 appear to evolve at a higher rate when considering mammal-wide data.

### Phylogenies based on full-length sequences

The phylogenetic signal in each protein sequence dataset was assessed by measuring the RF-distance between the tree resulting from that dataset and the reference tree, as well as manual inspection of differences in the topologies. There was no major difference in the phylogenetic trees created by maximum likelihood (ML) or Bayesian analysis (Figure 4). The Bayesian analysis performed slightly better by creating more accurate trees (smaller RF-distances to reference tree) from the 5 (ML = 170, Bayesian = 153) and 10 (ML = 122, Bayesian = 117) protein concatenations; however, the ML approach produced a slightly more accurate tree for the 14 (ML = 108, Bayesian = 110) protein concatenation. For this reason, all following analyses were performed using the ML approach, which also shows a higher computational efficiency. In all trees, all taxa were placed correctly at least at the family level, with two exceptions: First, in the tree based on the 5 protein concatenation, Galagonidae and Lorisidae remain unresolved. Second, while the general tendency is that the more proteins that are included in the concatenation, the more similar the tree is to the reference tree, there is one caveat for the 14 protein concatenation. The deepest relationship within Primates, namely the branching pattern between lorises and lemurs (Strepsirrhini), tarsiers (Tarsiiformes), and monkeys and apes (Simiiformes), is incorrectly resolved. However, in the phylogenies from the 5 and 10 protein concatenations it agrees with the reference tree (supplementary figure S2). In the trees of the 14 protein concatenation (ML and bayesian), Tarsiiformes form a clade with Strepsirrhini to the exclusion of Simiiformes (Figure 5) with a bootstrap value of 90 and a posterior probability of 1; however, current molecular and morphological evidence (Hartig et al. 2013; Morse et al. 2019; Seiffert et al. 2020; Kuderna et al. 2023) collectively provides compelling support for Tarsiiformes+Simiiformes to the exclusion of Strepsirhini.

**Figure 4:**
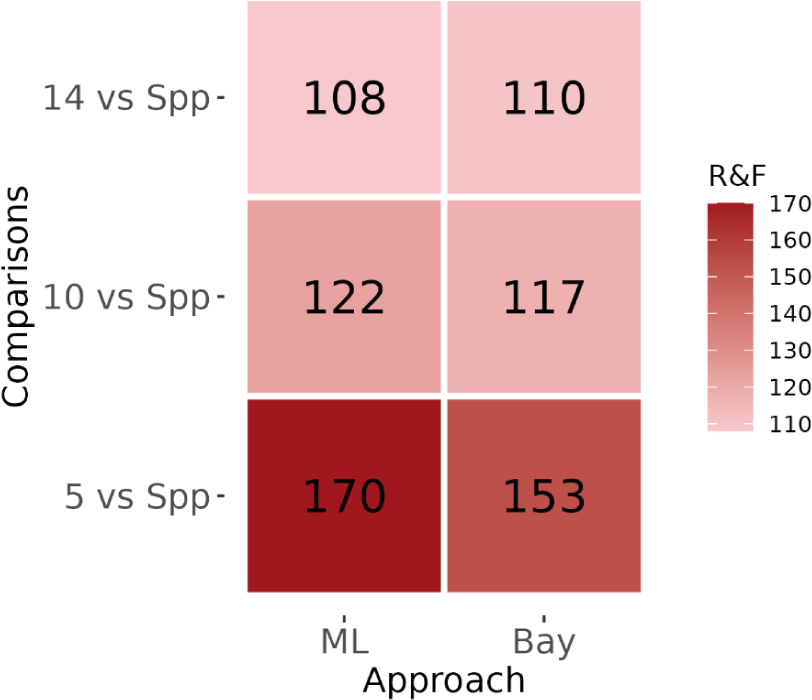
Robinson-Foulds distances between phylogenies based on 5, 10, and 14 protein concatenation and reference species tree for maximum likelihood and Bayesian approach. Spp - Reference species tree based on whole genome data (Kuderna et al. 2023), ML - maximum likelihood approach (IQ-TREE v. 1.6.12), Bay - Bayesian approach (MrBayes v.3.2.7a).

**Figure 5:**
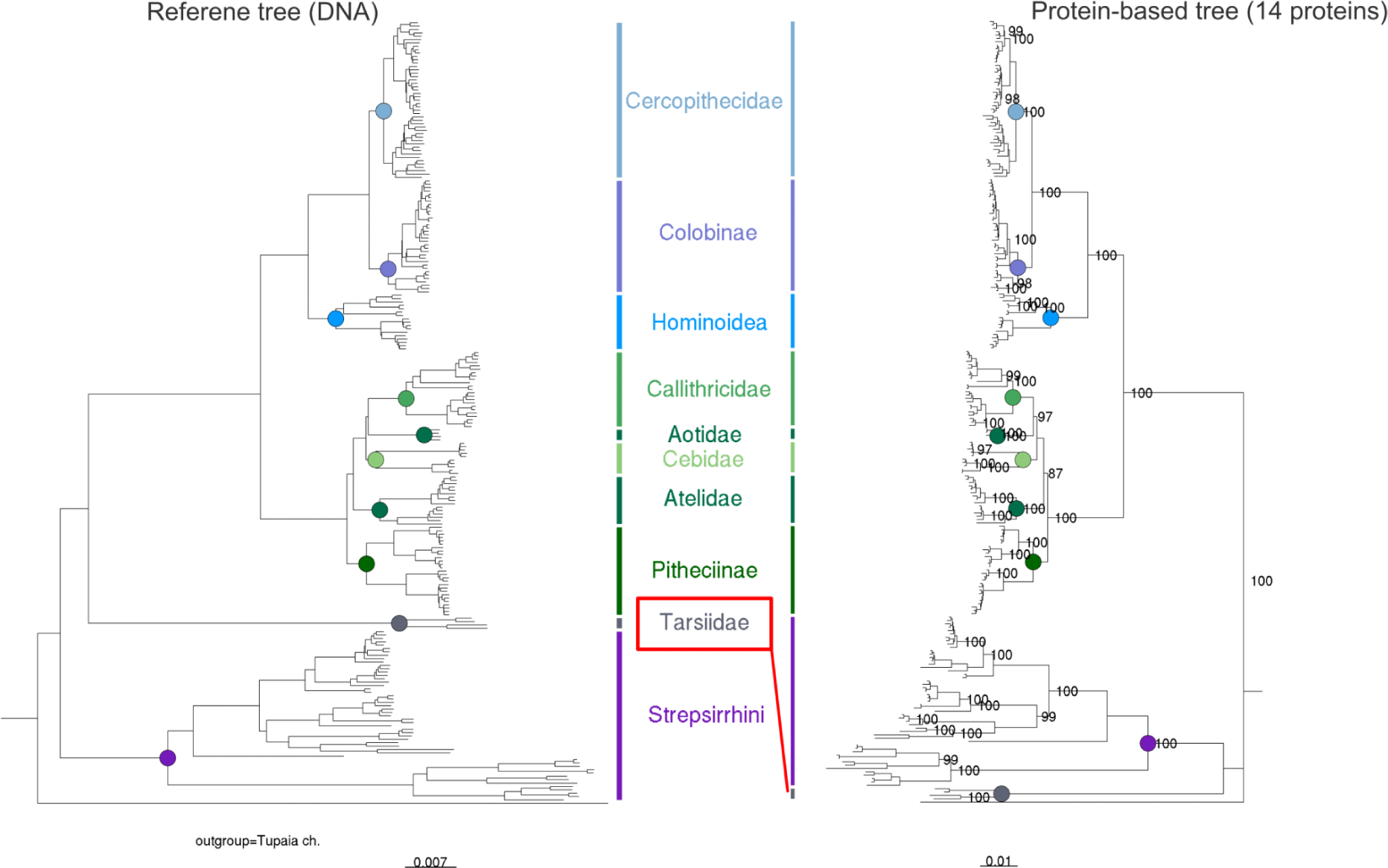
Species tree compared to phylogenetic tree based on 14 protein concatenation. The placement of families and even genera is largely in accordance. However Tarsiiformes (only family Tarsiidae) form a monophyly with Strepsirrhini (node of last common ancestor not visible because of short branch length), a placement that is nowadays widely rejected (Hartig et al. 2013; Morse et al. 2019; Seiffert et al. 2020; Kuderna et al. 2023). In the phylogenies based on concatenations of 5 and 10 proteins, which do not comprise collagens, Tarsiidae are placed as a sister group of Simiiformes in accordance with the reference tree (Kuderna et al. 2023) (supplementary figure S2).

In our case, the addition of four collagen genes to the dataset, resulting in the 14 protein concatenation, drove the misplacement of Tarsiiformes. We tested different combinations of collagens and non-collagen proteins of our dataset to see which genes in particular are driving this misplacement (supplementary table S3). A phylogeny based on only COL1A1, COL1A2, or COL17A1, respectively, places Tarsiiformes with Strepsirrhini. A phylogeny based on COL2A1 alone places them correctly with Simiiformes. If any of these individual collagens is combined with all 10 non-collagen genes into an 11 protein concatenation, the non-collagen genes drive the placement of tarsiers to the correct position, according to the species tree. However, if the 10 non-collagen genes are combined with COL1A2 and COL1A1 or COL17A1, this is sufficient to override the signal in the non-collagen proteins and place Tarsiiformes with Strepsirrhini.

### Phylogenies by fragmentation stage

Based on the modeled fragmentation stages, we created four MSAs with increasingly reduced sequence data and examined phylogenetic trees that were calculated from those MSAs. The protein concatenation corresponding to the fragmentation stage of “100 ka’’ had a total length of 3884 amino acids (Table 1). The phylogeny based on this data showed a RF-distance of 156 to the species tree (Figure 6), in contrast with the phylogeny based on the full-length 14 proteins (alignment length 9557 amino acids) which showed an RF-distance to the species tree of 108. All placements at family level and mostly genus level are in accordance with the reference tree, except for Tarsiiformes being grouped with Strepsirrhini (for more details see supplementary table S5). This is in line with the previous observation that the inclusion of COL1A2 and COL1A1 or COL17A1 can produce this result, even if just fragments of these protein sequences are included. Only five genera *(Cebuella*, *Lophocebus*, *Chlorocebus*, *Semnopithecus*, *Rhinopithecus*) are placed differently to the reference tree, but still within their correct family. Consequently, most differences that explain the RF-distance of 156 stem from different placements of species within their genus. The phylogeny based on the fragmentation stage “1-2 Ma” (MSA length 1139 aa) has a very similar distance to the species tree (158). Most nodes at family level are placed in accordance with all confidently resolved nodes of the reference tree, with some exceptions. Tarsiiformes are placed as an outgroup to both Strepsirhini and Simiiformes (bootstrap 100), a placement that is rejected by current molecular and morphological evidence (Hartig et al. 2013; Morse et al. 2019; Seiffert et al. 2020; Kuderna et al. 2023). In platyrrhines, Atelidae are placed as a sister clade to Callitrichidae (bootstrap 81), however, in the reference tree, it forms an outgroup to Cebidae, Callitrichidae and Aotidae. Moreover, in Lemuroidea, Galagidae are placed within the clade of Lorisidae (bootstrap 89), instead of being a sister taxon to Lorisidae, as in the reference tree. Some genera in the Cercopithecidae are placed slightly differently to the reference tree. In Hominidae, contrasting the reference tree, *Pan* and *Gorilla* are sister taxa, with *Homo* as an outgroup.

**Figure 6:**
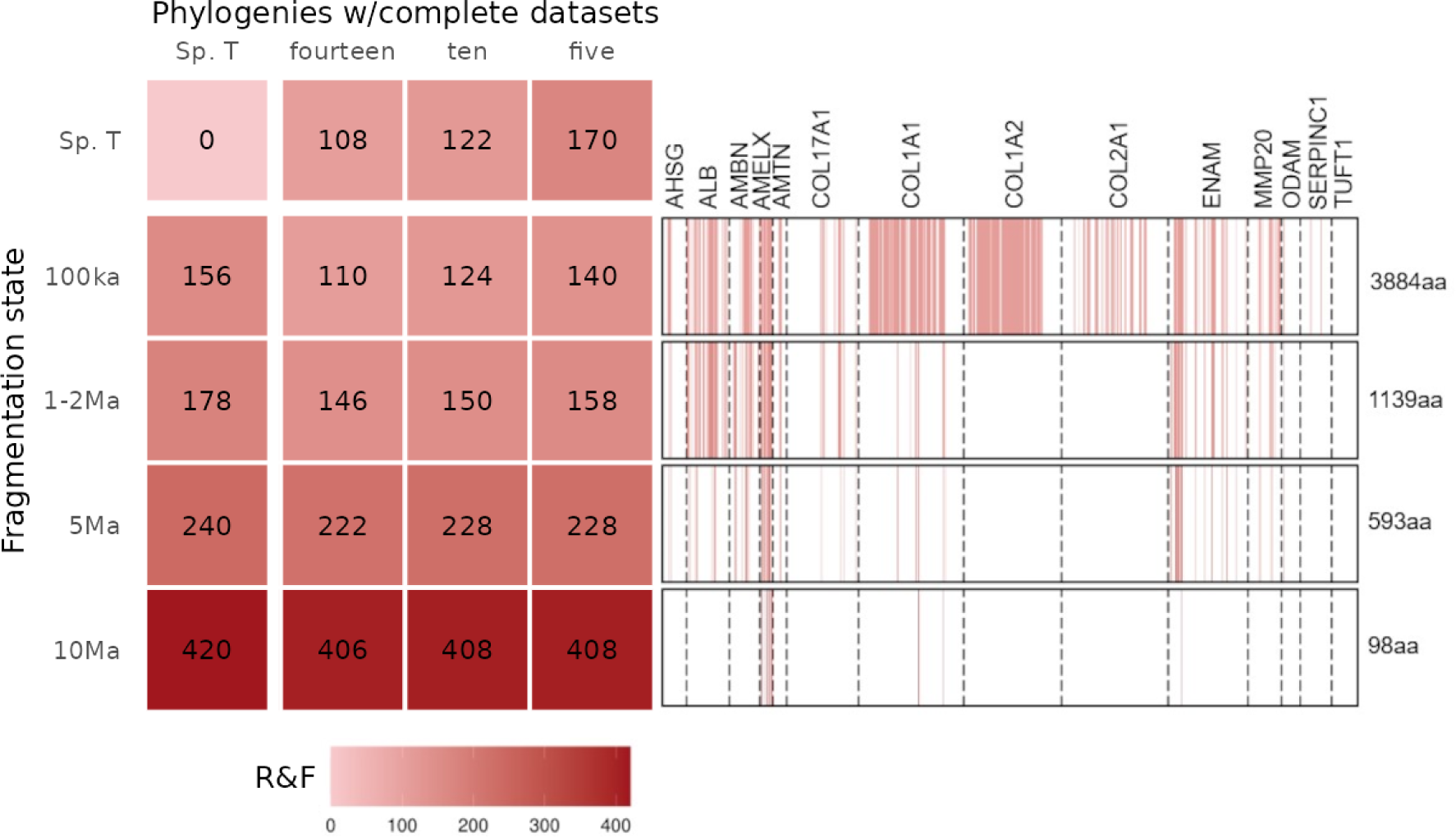
Robinson-Foulds distances between phylogenies based on simulated ancient data and reference tree. Spp - Reference tree based on whole genome data (Kuderna et al. 2023), “100 ka”, “1-2 Ma”, “5 Ma”, “10 Ma” represent different stages of fragmentation to which the amino acid MSA has been reduced prior to phylogenetic analysis.

**Table 1:** Details of multiple sequence alignments.

| Number of<br>proteins in<br>concate-<br>nation | Proteins | Length MSA<br>full sequence<br>(w/o signal) /<br>variable sites<br>/ parsimony<br>inform. sites<br>(aa) | Length<br>MSA<br>fragmented<br>“100 ka” | Length<br>MSA<br>fragmented<br>“1-2 Ma” | Length<br>MSA<br>fragmented<br>“5 Ma” | Length<br>MSA<br>fragmented<br>“10 Ma” | Length of<br>each<br>“variable” /<br>“conserved<br>“ MSA |
| --- | --- | --- | --- | --- | --- | --- | --- |
| 5 | AMBN,<br>AMELX,<br>AMTN, ENAM,<br>MMP20 | 2339 / 1443 /<br>1225 | NA | NA | NA | NA | shannon:<br>826 / 1513<br>r4s:<br>878 / 1461 |
| 10 | AMBN,<br>AMELX,<br>AMTN, ENAM,<br>MMP20, | 4327 / 2547 /<br>2118 | NA | NA | NA | NA | shannon:<br>1535 / 2792<br>r4s:<br>1557 / 2770 |
|  | AHSG, ALB,<br>ODAM,<br>SERPINC1,<br>TUFT1 |  |  |  |  |  |  |
| 14 | AMBN,<br>AMELX,<br>AMTN, ENAM,<br>MMP20,<br>AHSG, ALB,<br>ODAM,<br>SERPINC1,<br>TUFT1,<br>COL1A1,<br>COL1A2,<br>COL2A1,<br>COL17A1 | 9557 / 3636 /<br>3050 | 3884 | 1139 | 593 | 98 | shannon:<br>3632 / 5925<br>r4s:<br>2516 / 7041 |

The tree based on data of the fragmentation stage “5 Ma’’ (alignment length 593 amino acids) has an RF-distance to the species tree of 240. In catarrhines, all family level relationships still agree with the reference tree, but there are some inconsistencies within families compared to the reference tree. Within Hominidae, *Pan* and *Gorilla* are sister taxa; all cercopithecid species are still placed in the correct family but the placement of most genera differs from species tree, often with bootstrap values below 50. In platyrrhines, correct resolution of nodes at family level is widely lost. In contrast to the species tree, Pitheciidae are not recovered as monophyletic. Some genera in Callitrichidae are no longer monophyletic, and part of Cebidae is placed with them. The placement of nodes at family level in Lemuridae is in accordance with the species tree, except for *Varecia* no longer being an outgroup to Lemuridae, but placed within Lemuridae. Similarly, *Daubentonia madagascariensis* is no longer the outgroup to Lemuriformes, but placed within them (bootstrap 92), with Lemuridae forming the basal-most branch of this clade (bootstrap 100). Tarsiiformes form a clade with Strepsirhini (bootstrap 93).

At the fragmentation stage “10 Ma” (alignment length 98 amino acids), the phylogeny is largely unresolved, with a RF-distance to the reference tree of 420 and most bootstrap values far below 50. Lorisiformes are correctly separated as their own taxon. All species of Malagasy primates (i.e. the chiromyiform *Daubentonia* and Lemuriformes) form a clade, but some platyrrhines (e.g. *Sapajus apella*, *Callithrix* spp., *Callimico goeldii*) and some catarrhines (e.g. *Rhinopithecus bieti*, *Nasalis larvatus*) are placed within them. Most catarrhine species still group together, but platyrrhines no longer form a clade. The four Tarsiiformes species are monophyletic and placed with low confidence within the other groups.

### Case studies

Often, paleoproteomic studies will only aim to place a few closely related specimens at a time into a framework of mostly complete reference protein sequences. To simulate such a scenario, we created two cases in which protein sequence data from a group with fairly well-known taxonomic placement is fragmented and aligned to reference sequences that are at full-length. In the first case, sequence data of two Neanderthals (individuals Vindija 33.19 and Altai) and one Denisovan (Denisova3 individual) that were fragmented to the degradation stage “100 ka” was aligned to a reference MSA of 14 enamel-related proteins from Hominoidea. The phylogenetic placement is in accordance with the reference tree at genus level, but not at species level (supplementary figure S5A). The Neanderthal and Denisovan individuals place within the *Homo sapiens* clade with low branch support, instead of forming a sister clade. In the same tree, the two species with the youngest splits, *Pongo abelii* and *Pongo pygmaeus* do not form two distinct clades, nor do *Gorilla gorilla* and *Gorilla beringei*, despite being based on full-length protein data. These results align with previous observations on the limitations of paleoproteomic data to resolve phylogenies in Hominidae at species-level resolution (Welker et al. 2019; Welker et al. 2020; Madupe et al. 2023). In particular, if several individuals per clade are examined using such limited sequence data, the differences between individuals can override the signal that separates the species (Madupe et al. 2023). The example of simulated ancient chimpanzee data of a fragmentation stage of “5 Ma’’ produces a different result (Supplementary figure S5B). While in the reference *Pongo* and *Gorilla* cannot be resolved at species level, the three *Pan troglodytes* individuals are placed confidently as a sister taxon to *Pan paniscus* (bootstrap 100). Divergence times have been estimated using MCMCtree for each node of a simplified version (one individual per species) of the “Chimpanzee case” tree (Figure 7, supplementary table S6, supplementary figure S6). Most divergence time estimates derived from the protein-based data of the “Chimpanzee case” are younger than those of the genome-based reference (a mean of 21% younger). Only the divergence time estimate of the split between the *Pan troglodytes* (fragmented protein sequences) and *Pan paniscus* (complete protein sequences) is, with a mean of 3.58 Ma, 50% older than that of the reference (Kuderna et al. 2023). The 95% highest posterior density intervals (95% HPD) of all internal nodes overlap between the reference and our case study, with the exception of the split between Ponginae and Homininae (12.24 - 16.42 Ma in our case study, 18.58 - 22.19 Ma in reference). While the boundaries of 95% HPD of this case study are rather close to the effective prior’s 95% HPD boundaries (12.25 - 15.26 Ma), the boundaries of 95% HPD of the reference are completely outside their effective prior 95% HPD bounds (12.26 - 14.58 Ma). This may be a result of genomic data contributing signals that produce a posterior distribution with significantly older ages. These signals, however, may not be reflected in the protein sequences of the enamelome, which comprise a much smaller dataset (MSA length 9557 amino acids), compared to the genomic data (676 kilo base pair, kbp, long MSA of ultra conserved elements). In addition, the analysis presented here only estimated divergence dates for the superfamily Hominoidea (apes and humans) whereas Kuderna et al. (2023) calculated divergence dates for all primates, which allowed the use of additional fossil calibrations proposed by de Vries and Beck (2023).

**Figure 7:**
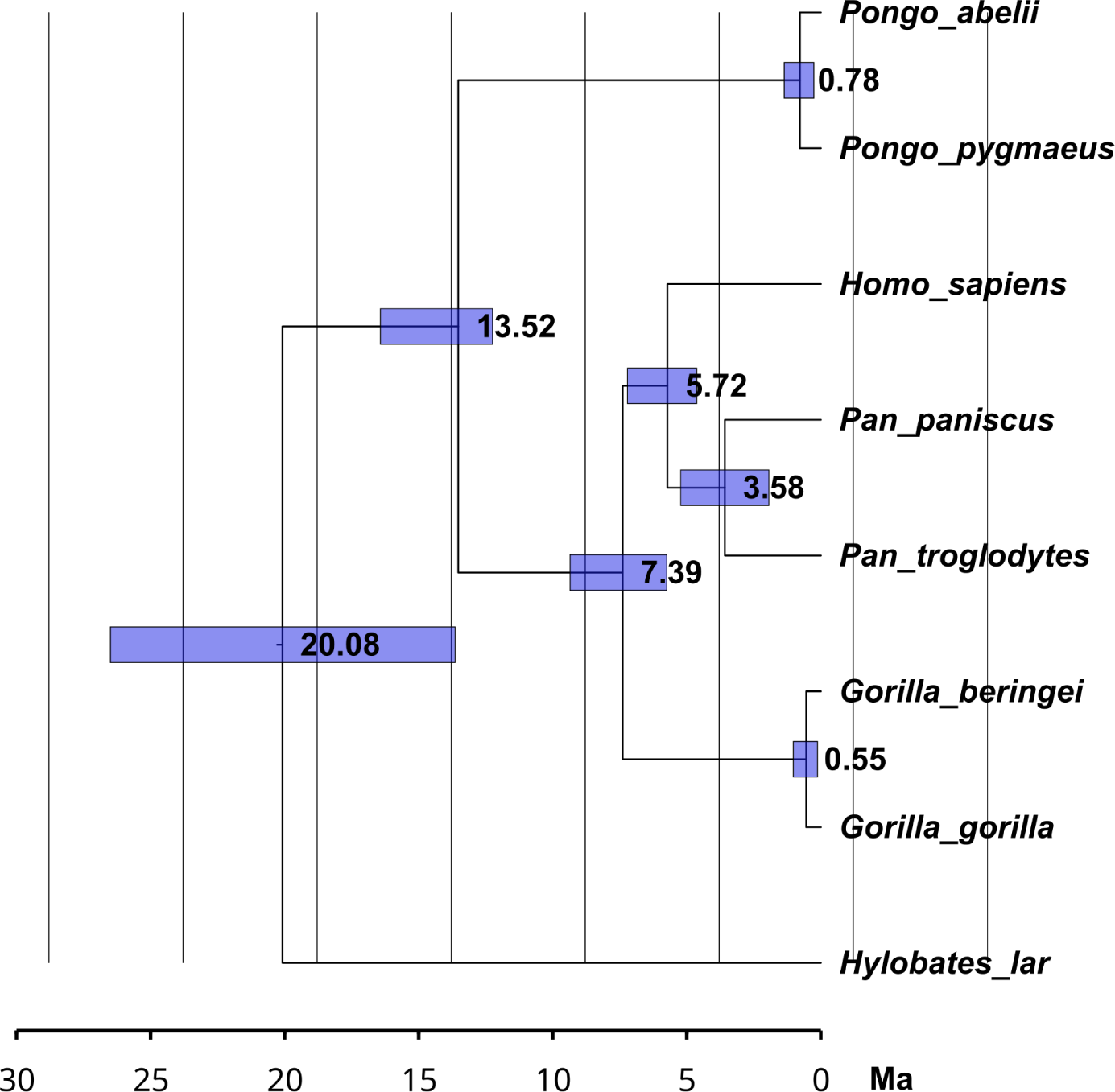
Phylogeny with divergence time estimates for “Chimpanzee case” simulation: The X-axis represents million years before present. Blue bars display 95% HPD of all 10 MCMCtree runs combined. All 95% HPD are overlapping with the ones of the reference tree (Kuderna et al. 2023), including the simulated ancient *Pan troglodytes* sequence. Only the 95% HPD of the split between Ponginae and Homininae covers a younger time range and does not overlap with that of the reference tree (12.24 - 16.42 Ma in our case study, 18.58 - 22.19 Ma in reference).

### Phylogenies by amino acid conservation

To quantify the contribution of the variable sites in each MSA to the correct resolution of the according tree, we divided all concatenated MSAs into two MSAs, one consisting of all variable sites, the other of all conserved sites. To divide between variable and conserved, the MSA of each protein was normalized to a mean of 1 in their Shannon entropy values or Rate4Site scores. They were then divided into sites above this value, i.e. the “variable” sites, and below, i.e. the “conserved” sites (supplementary figure S3). These sites from all proteins were concatenated and used for phylogenetic analysis with ML. Phylogenies based only on variable sites identified by Rate4Site (Figure 8, “R4S variable”) are just as similar to the species tree as in the case of phylogenies based on the full-length protein sequences (Figure 4, both have a RF-distance of 108 to the reference tree). When using Shannon entropy to define conserved and variable sites, phylogenies based on the variable sites (Figure 8, “Shannon variable”) are more similar to the species tree than those based on the more conserved sites (“Shannon conserved”). Nonetheless, all phylogenies based on variable sites that were identified using Rate4Site (“R4S variable”) are more similar to the reference tree than those based on variable sites that were identified using Shannon entropy (“Shannon variable”).

**Figure 8:**
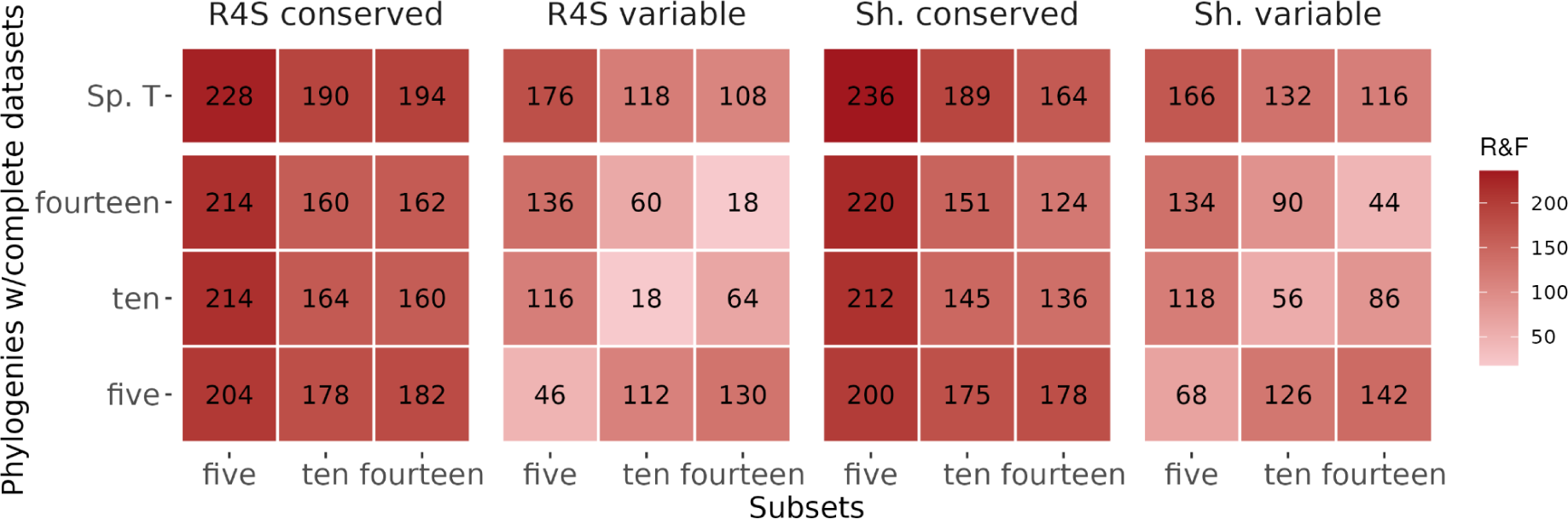
Robinson-Foulds distances between phylogenies based on highly conserved or variable data and reference species tree. The phylogenies based on variable sites show comparably small RF-distances to the reference tree (Kuderna et al. 2023) (see Figure 4; 5 protein tree: 170, 10 protein tree: 122, 14 protein tree: 108). Especially, Rate4Site identifies the sites that drive the topology to resemble the reference. Phylogenies based on the conserved sets of sites tend to show larger RF-distances.

### ERC scores

The enamel proteome is functionally linked, in particular because all proteins are expressed only during amelogenesis (formation of enamel) during a very short phase of an individual’s development (Castiblanco et al. 2015). In addition, the amelogenins (AMELX and AMELY) share sequence homology, as part of the same gene family. The same applies to all collagens. Correlated evolutionary rates and sequence similarity between proteins are factors to take into account when creating a molecular phylogeny based on such a limited protein sequence dataset. We estimated the degree of evolutionary co-variation of the set of 14 genes using Evolutionary Rate Covariation (ERC) analysis (Clark et al. 2012). ERC returns pairwise correlation coefficients of the branch-specific evolutionary rates of a set of genes (Figure 8). Permutation testing indicated that all pairwise values between amelogenesis proteins are significantly elevated (p-value < 0.0001). There is particularly elevated covariation (Fisher-transformed value > 3) in the evolutionary rates of all pairs between COL1A1, COL1A2, and COL2A1. COL17A, the most divergent of all of the collagens and the only one known to have a function in enamel formation, displays lower covariation with the other collagens. AMBN, AMTN, ENAM, and ODAM are located on a syntenic block (e.g. *Homo sapiens*, chr 4; *Pan troglodytes*, chr 4; *Microcebus murinus*, chr 26, *Mus musculus*, chr 5). The elevated and significant ERC values between them may reflect the functional and spatial proximity of these genes. One of the genes with the highest pairwise correlation values is MMP20, a gene that encodes for an enzyme that cleaves AMBN, AMELX, AMELY, and ENAM during amelogenesis (Gasse et al. 2017). All genes encoding those cleavage targets display elevated values of covariation of evolutionary rates. AMTN is another gene that displays a higher correlation in evolutionary rates with the aforementioned group, but little is known about its interactions and function. The most striking degree of covariation (Fisher-transformed value = 19.2) can be observed between AMELX (chromosome X) and AMELY (chromosome Y). For being encoded on the sex chromosomes they can be considered paralogs. In humans, AMELY lies outside of the pseudoautosomal region. However, it is known that both are expressed, if a Y chromosome is present, and AMELX and AMELY seem to fulfill the same function (Haruyama et al. 2011; Parker et al. 2019).

## Discussion

In this study, we estimated the degree of sequence conservation, evolutionary rate, and phylogenetic signal of protein sequences that are associated with the primate enamel proteome. The analyses put an emphasis on evaluating these measures from a perspective of experimental feasibility, since ancient peptide data is highly fragmentary and diagenetically altered. The process of degradation was simulated for different stages of fragmentation, which were anticipated from experimental data. Given the limited amount of experimental data and the over-representation of samples younger than 2 Ma, it has not yet been possible to statistically assess the patterns of *post mortem* sequence degradation, but will hopefully be so in the future as more ancient enamel peptide sequences are published. However, patterns are already visible, e.g. the deep time sequence survival of N-terminal peptides of ENAM (Figure 3) or that of the N-terminal region of AMELX and its C-terminal proline rich region (Figure 3). When simulating fragmented data and subsequently performing phylogenetic inference, most families were in accordance with the reference tree up to a fragmentation stage similar to published samples of an age of 1-2 Ma from temperate to tropical regions. This does not exclude that sequences with a higher degree of fragmentation (stage “5 Ma”) could be placed correctly in a phylogeny, as the *Pan troglodytes* sequences in our case study were rather highly fragmented (stage “5 Ma”) and still correctly placed. Also, the estimation of its mean divergence time was 50% older than the reference, but still had an 95% HPD that completely overlapped with that of the reference (Kuderna et al. 2023). In this particular case study, the divergence time estimates were generally lower and closer to the effective prior distribution than those of the reference. This may be a reflection of the relatively high degree of sequence conservation of the enamelome (Bartlett et al. 2006; Al-Hashimi et al. 2009; Haruyama et al. 2011; Castiblanco et al. 2015; Gil-Bona and Bidlack 2020; Warinner et al. 2022). Moreover the overall short sequence length of all enamelome protein sequences combined is significantly shorter than that of the genomic data as used in the reference (9557 aa enamelome MSA vs. 676 kbp DNA reference-MSA). To fully understand the implications of using the enamelome for divergence time estimates, more comparative studies are needed.

Regarding tree topology, variable sites within these protein sequences were clearly more phylogenetically informative than the more conserved sites, especially if the category “variable” is determined using Rate4Site (Figure 8). This falls within the expectation, since Rate4Site accounts for amino acid replacement models and phylogenetic relationships between the input sequences. In contrast to the utility of ultra conserved elements in genomic data, the more conserved sites in our protein data, were not able to produce more reliable phylogenetic trees. Most likely because a large part of the conserved sites had no variability at all (Table 1). However, variable sites do not always fall within those regions that could be experimentally recovered, e.g. in AHSG, AMTN, COL17A, MMP20, ODAM or TUFT1 (Figure 3). It may be possible to adapt protocols for peptide isolation from tooth enamel. Some progress has been achieved recently by fractionating the sample in order to recover more fragments of different hydrophobicity (Madupe et al. 2023). Identifying variable sites in collagens may also be of interest for optimizing protocols for the application of ZooMS (Buckley et al. 2009; Naihui et al. 2021). The general pattern of conservation of individual sites can also be observed across mammals (supplementary figure S4). Some cases distinguish primates from the general trend in mammals, e.g. the 32 kDa fragment of ENAM appears particularly conserved in primates. In fact, in this region, signals of positive selection have been reported in primates (Al-Hashimi et al. 2009). This indicates that the degree of sequence conservation might not be consistent across clades.

Not only clades, but also single proteins can differ from each other in their evolutionary history (Mu et al. 2021). This may be a possible explanation for the case of tarsiers, which were placed in profoundly different locations within the phylogenetic tree depending on whether or not collagens were included in the dataset (Figure 5, supplementary table S3). However, we cannot rule out other causes, such as model misspecification, i.e. the collagen sequences may have evolved in a way that cannot be appropriately modeled by the phylogenetic model used in our analysis. This case example highlights a potential pitfall of paleoproteomics when used for phylogenetic analysis. The specimen of interest might be placed with a reasonably high confidence in a phylogeny based on concatenated protein sequences, as for example tarsiers were placed as “Prosimii” with Stresirrhini at the “100 ka” fragmentation stage. Still, as this example demonstrates, such a placement could be in conflict with current evidence, and, in contrast to this case, there may be no genomic data for the ancient sample that can be used to test the accuracy of the proteomic data.

Evolution of a set of proteins from a specialized tissue may be tightly linked due to the constraints of morphology and function of this tissue. Our example of tarsiers underlines why working with such a small tissue-specific set of biological sequences should be accompanied by morphometric and histological expertise. For example, compared to Simiiformes, Tarsiiformes and Strepsirrhini share the traits of thinner tooth enamel (Shellis et al. 1998), and similar enamel microstructure (Maas and Dumont 1999). Both may be reflected in a similar genetic basis, e.g. as an ancestral trait or as a result of convergent evolution.

Beyond the morpho-functional constraints, the relationships between such a small set of genes can be further entangled, as this is the case for AMBN, AMTN, ENAM, and ODAM, which are located in close proximity to each other on the same chromosome in most mammals (e.g. *Homo sapiens*, chr 4). This has also been reflected in significantly higher ERC values in our analysis (Figure 9) and partly observed in another study that could associate evolutionary rates of ENAM and ODAM to enamel thickness (Mu et al. 2021). A third aspect of covariation and possible codependence of this set of typically studied genes is high sequence similarity between some of them. For instance, all collagens in this study share 38-72% sequence identity among each other in humans (aligning UniProt entries P02452, P08123, P02458, and Q9UMD9). We do not have sufficient genomic data to include the Y-chromosomal AMELX paralog, AMELY, into our analyses based on predicted protein sequences. It is known to share around 88.5% sequence similarity with AMELX in humans (aligning UniProt entries Q99217-3 and Q99218-2) and it showed by far the highest degree of covariation in the ERC analysis. In other mammals, signs of gene conversion between AMELX and AMELY have been reported, indicating that these two genes and their protein sequences are not acting as independent *loci* (Janečka et al. 2018; Kawasaki et al. 2020). The dependencies that exist within this small proteome challenge the practice of concatenating them into a single, long MSA to address phylogenetic questions, because an overrepresented set of dependent *loci* might skew the outcome towards their shared evolutionary history. The difference between single-gene trees and species trees in the context of the enamel proteome has been demonstrated and discussed in previous publications (Welker et al. 2019; Welker et al. 2020), in which phylogenetic inference based on concatenated MSAs delivered results that were more consistent with the recognized species tree for the verifiable extant reference taxa. Most paleoproteomic studies have had and probably will have their main interest in the evolution of species, rather than single genes. Alternatives to the simple concatenation of DNA or protein sequences from single genes exist (Douglas et al. 2022), although they need more comparative assessment for their utility in paleoproteomics (Madupe et al., 2023). However, it still condenses to making an adequate choice of proteins to be considered, which is challenging because the set of proteins is small and the amount of amino acid sites that can be isolated and sequenced is typically in the range of 450 -1000 (Cappellini et al. 2019; Welker et al. 2019; Welker et al. 2020; Madupe et al. 2023).

**Figure 9:**
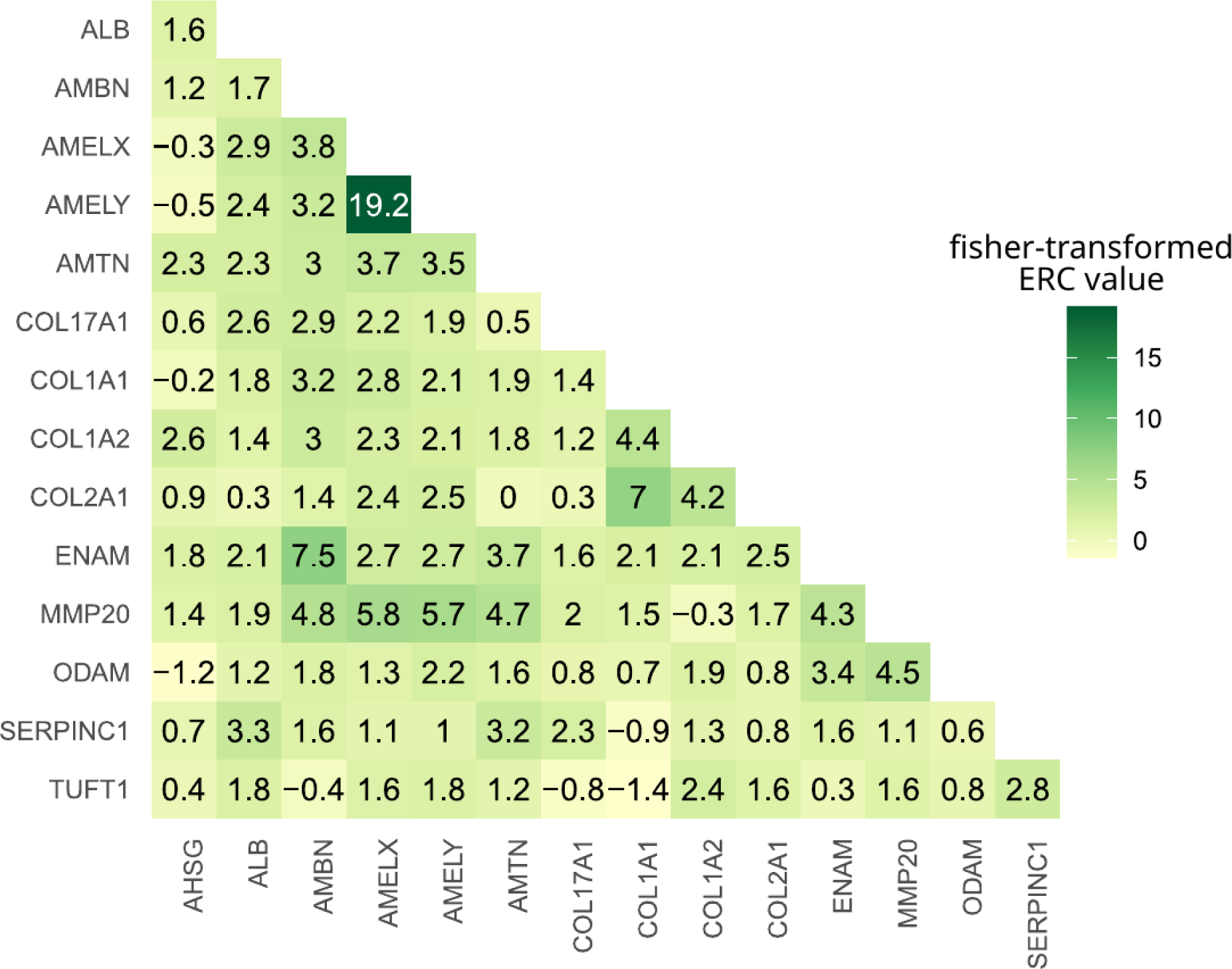
Fisher-transformed ERC values between the 14 proteins of this study. The strongest correlation of evolutionary rates was measured between AMELX and AMELY. Elevated ERC values can also be observed between the gene that encodes the enzyme MMP20 and its cleavage targets AMBN, AMELX, AMELY, and ENAM (Gasse et al. 2017). Also between the non-enamel collagens COL1A1, COL1A2, and COL2A1, elevated ERC values can be observed.

Consequently, for any paleoproteomic study in organismal evolution, we consider it advisable to perform a similar analysis of sequence conservation and phylogenetic signal of all available proteins of the taxonomic group and tissue of interest.

## Conclusion

Our set of tooth enamel-associated proteins revealed sites of higher and lower sequence conservation within each protein, but collagens evolve at overall slower rates than non-collagenous enamel proteins. Phylogenies that are based on the more variable sites of all proteins combined tend to be closer to the reference species tree. If the protein sequences are fragmented to simulate paleoproteomic experimental data, the placement of the vast majority of families and genera is in accordance with the reference species tree up to a fragmentation stage that corresponds to most published samples (1-2 Ma old). Further fragmentation leads to incorrect placement of nodes at family level, but may still produce reliable results if only one individual has fragmented sequence data. With the case of collagens and tarsiers, we could demonstrate that the inclusion or exclusion of information from single proteins can sometimes change the placement of single clades dramatically. Our results demonstrate that each protein and taxon have their own evolutionary history, which can influence the results of analyses with such limited data, as it is the case in paleoproteomics. Therefore, for the phylogenetic application of paleoproteomic data, we suggest exploring the relevant protein sequences of the closest related extant species.

## Supporting information

Supplementary information including SUPPLEMENTARY FIGURES and TABLES

List of all individuals

Sites that could not be translated, manually marked with "?" or "X"

MSAs of all 14 proteins ("BLOCK1") and MSAs of ancient sequences ("BLOCK2")

Sites referring to columns in MSAs of supplementary file 3 ("BLOCK1") to model sequence degradation

## Data availability

The mass spectrometry proteomics data have been deposited to the ProteomeXchange Consortium (Deutsch et al. 2017) via the PRIDE (Perez-Riverol et al. 2022) partner repository with the dataset identifier PXD048972. Additional supplementary data to the supplementary files associated with this publication, is deposited at Zenodo under the doi 10.5281/zenodo.10637110. All code is available under github.com/RicardoFong/primate_enamelome.

## Acknowledgements

J.K. is supported by the European Union’s Horizon 2020 research and innovation program under the Marie Sklodowska-Curie “PUSHH” training network, grant agreement No. 861389. T.M.B. is supported by funding from the European Research Council (ERC) under the European Union’s Horizon 2020 research and innovation programme (grant agreement No. 864203), PID2021-126004NB-100 (MICIIN/FEDER, UE) and Secretaria d’Universitats i Recerca and CERCA Programme del Departament d’Economia i Coneixement de la Generalitat de Catalunya (GRC 2021 SGR 00177). This work is part of R+D+I projects PID2020-116908GB-I00 and PID2020-117289GB-I00, funded by the Agencia Estatal de Investigación of the Spanish Ministerio de Ciencia e Innovación (MCIN/AEI/10.13039/501100011033/). Research has also been supported by the Agència de Gestió d’Ajuts Universitaris i de Recerca of the Generalitat de Catalunya (2001 SGR 00620). J.D.O. was supported by the ”la Caixa” Foundation (ID 100010434) and the European Union’s Horizon 2020 research and innovation program under the Marie Skłodowska-Curie grant agreement No 847648 (LCF/BQ/PI20/11760004), and the National Sciences Research Council of Canada (RGPIN-2023-04399, DGECR-2023-00272). O.C. wish to acknowledge the “Juan de la Cierva Formación” program (ref. JDC2022-048590-I), funded by the Agencia Estatal de Investigación of the Spanish Ministry of Science and Innovation (MCIU/AEI/10.13039/501100011033) and the European Union “NextGenerationEU”/PRTR” program. R.M.D.B.’s research was supported by NERC Standard Grant ‘Rise of the continent of the monkeys’ (NE/T000341/1). We thank Claudia Fontsere, Alejandro Valenzuela, Mareike Janiak, Frido Welker, and Dorien de Vries for their advice during project planning and data analysis.

## Author contributions

Conceptualization: E.L., J.K., and R.F.Z. Methodology and analyses: R.F.Z. and J.K. with assistance and guidance from all co-authors. Evolutionary Rate Covariation scores were calculated by J.L. and N.C. Writing: J.K. and R.F.Z. with input from all co-authors.

## Supplementary information

See separate document

