## Supplementary information including SUPPLEMENTARY FIGURES and TABLES for "Phylogenetic signal in primate tooth enamel proteins and its relevance for paleoproteomics"

### 1.Dataset generation

All mentioned code that is not explicitly explained here, is available under github.com/RicardoFong/primate_enamelome.

#### 1.1 Prediction of Protein sequences

The dataset consisted of 722 individuals of 233 species, spanning the 16 families of the primate order. Identifiers and species are available in supplementary file1.

Not every species has a reference assembly, so that the individuals of some species were mapped to one of their closest relatives "reference species". If not specified otherwise, the assembly for a species is the same as in Kuderna et al. (2023). The mapping went as follows:

Reference species: Aotus_nancymaae

Species mapped:

- Aotus_azarae
- Aotus_griseimembra
- Aotus_trivirgatus
- Aotus_vociferans

Reference species: Atele_fusciceps

Species mapped:

- Alouatta_belzebul
- Alouatta_caraya
- Alouatta_discolor
- Alouatta_juara
- Alouatta_macconnelli
- Alouatta_nigerrima
- Alouatta_palliata
- Alouatta_puruensis
- Alouatta_seniculus
- Ateles_belzebuth
- Ateles_chamek
- Ateles_geoffroyi
- Ateles_marginatus
- Ateles_paniscus
- Lagothrix_lagothricha

Reference species: Callithrix_jacchus

Species mapped:

- Callibella_humilis
- Callimico_goeldii
- Callithrix_geoffroyi
- Callithrix_jacchus
- Callithrix_kuhlii
- Cebuella_niveiventris
- Cebuella_pygmaea

Leontopithecus_chrysomelas

Leontopithecus_rosalia

Mico_argentatus

Mico_humeralifer

Mico_spnv

Reference species: Cebus_albifrons

Species mapped:

Cebus_albifrons

Cebus_olivaceus

Cebus_unicolor

Saimiri_cassiquiarensis

Saimiri_macrodon

Saimiri_oerstedii

Saimiri_sciureus

Saimiri_ustus

Sapajus_macrocephalus

Reference species: Cercocebus_atys

Species mapped:

Cercocebus_chrysogaster

Cercocebus_lunulatus

Cercocebus_torquatus

Reference species: Cercopithecus_mitis

Species mapped:

Cercopithecus_ascanius

Cercopithecus_cephus

Cercopithecus_diana

Cercopithecus_hamlyni

Cercopithecus_lowei

Cercopithecus_mitis

Cercopithecus_mona

Cercopithecus_neglectus

Cercopithecus_nictitans

Cercopithecus_petaurista

Cercopithecus_pogonias

Cercopithecus_roloway

Miopithecus_ogouensis

Reference species: Chlorocebus_aethiops

Species mapped:

Allenopithecus_nigroviridis

Allochrocebus_lhoesti

Allochrocebus_preussi

Allochrocebus_solatus

Chlorocebus_pygerythrus

Reference species: Colobus_guereza

Species mapped:

Colobus_angolensis

Colobus_guereza

Colobus_polykomos

Reference species: Daubentonia_madagascariensis

Species mapped:

Daubentonia_madagascariensis

Reference species: Erythrocebus_patas

Species mapped:

Erythrocebus_patas

Reference species: Galago_moholi

Species mapped:

Galagoides_demidoff

Galago_moholi

Galago_senegalensis

Reference species: Gorilla_gorilla_gorilla

Species mapped:

Gorilla_beringei

Gorilla_gorilla

Reference species: Homo_sapiens (hg19)

Species mapped:

Homo_sapiens

Neanderthal

Denisovan

Gorilla_beringei (partly, see methods)

Gorilla_gorilla (partly, see methods)

Pan_troglodytes (partly, see methods)

Pan_paniscus (partly, see methods)

Pongo_abelii (partly, see methods)

Pongo_pygmaeus (partly, see methods)

Reference species: Lemur_catta

Species mapped:

Eulemur_albifrons

Eulemur_collaris

Eulemur_coronatus

Eulemur_flavifrons

Eulemur_fulvus

Eulemur_macaco

Eulemur_mongoz

Eulemur_rubriventer

Eulemur_rufus

Eulemur_sanfordi

Hapalemur_alaotrensis

Hapalemur_gilberti

Hapalemur_griseus

Hapalemur_meridionalis

Hapalemur_occidentalis

Lemur_catta

Prolemur_simus

Varecia_rubra

Varecia_variegata

Reference species: Loris_tardigradus

Species mapped:

Loris_lydekkerianus

Loris_tardigradus

Reference species: Macaca_mulatta

Species mapped:

Macaca_arctoides

Macaca_assamensis

Macaca_cyclopis

Macaca_fascicularis

Macaca_fuscata

Macaca_leonina

Macaca_maura

Macaca_mulatta

Macaca_nemestrina

Macaca_nigra

Macaca_radiata

Macaca_siberu

Macaca_silenus

Macaca_thibetana

Macaca_tonkeana

Reference species: Mandrillus_sphinx

Species mapped:

Lophocebus_aterrimus

Mandrillus_leucophaeus

Mandrillus_sphinx

Reference species: Microcebus_murinus

Species mapped:

Cheirogaleus_major

Cheirogaleus_medius

Lepilemur_ankaranensis

Lepilemur_dorsalis

Lepilemur_ruficaudatus

Lepilemur_septentrionalis

Microcebus_murinus

Mirza_zaza

Reference species: Nomascus_leucogenys

Species mapped:

Hoolock_hoolock

Hylobates_abbotti

Hylobates_agilis

Hylobates_klossii

Hylobates_lar

Hylobates_muelleri

Nomascus_annamensis

Nomascus_concolor

Nomascus_gabriellae

Nomascus_leucogenys

Nomascus_siki

Reference species: Nycticebus_pygmaeus

Species mapped:

Arctocebus_calabarensis

Nycticebus_bengalensis

Nycticebus_coucang

Nycticebus_pygmaeus

Perodicticus_ibeanus

Perodicticus_potto

Reference species: Otolemur_garnettii

Species mapped:

Otolemur_crassicaudatus

Otolemur_garnettii

Reference species: Pan_troglodytes

Species mapped:

Pan_troglodytes

Reference species: Papio_anubis

Species mapped:

Papio_anubis

Papio_cynocephalus

Reference species: Pithecia_pithecia

Species mapped:

Cacajao_ayresi

Cacajao_calvus

Cacajao_hosomi

Cacajao_melanocephalus

Cheracebus_lucifer

Cheracebus_lugens

Cheracebus_regulus

Cheracebus_torquatus

Chiropotes_albinasus

Chiropotes_israelita

Chiropotes_sagulatus

Pithecia_albicans

Pithecia_chrysocephala

Pithecia_hirsuta

Pithecia_mittermeieri

Pithecia_pissinattii

Pithecia_vanzolinii

Plecturocebus_bernhardi

Plecturocebus_brunneus

Plecturocebus_caligatus

Plecturocebus_cinerascens

Plecturocebus_cupreus

Plecturocebus_dubius

Plecturocebus_grovesi

Plecturocebus_hoffmannsi

Plecturocebus_miltoni

Plecturocebus_moloch

Reference species: Pongo_abelii

Species mapped:

Pongo_abelii

Pongo_pygmaeus

Reference species: Propithecus_coquereli

Species mapped:

Avahi_laniger

Avahi_peyrierasi

Indri_indri

Propithecus_coquereli

Propithecus_coronatus

Propithecus_diadema

Propithecus_edwardsi

Propithecus_perrieri

Propithecus_tattersalli

Propithecus_verreauxi

Reference species: Macaca mulatta

Species mapped:

Papio_anubis

Papio_cynocephalus

Papio_hamadryas

Papio_kindae

Papio_papio

Papio_ursinus

Reference species: Rhinopithecus_roxellana

Species mapped:

Nasalis_larvatus

Piliocolobus_badius

Piliocolobus_gordonorum

Piliocolobus_kirkii

Piliocolobus_tephrosceles

Presbytis_comata

Presbytis_mitrata

Pygathrix_cinerea

Pygathrix_nemaeus

Pygathrix_nigripes

Rhinopithecus_bieti

Rhinopithecus_roxellana

Semnopithecus_entellus

Semnopithecus_hypoleucos

Semnopithecus_johnii

Semnopithecus_priam

Semnopithecus_schistaceus

Semnopithecus_vetulus

Trachypithecus_auratus

Trachypithecus_crepusculus

Trachypithecus_cristatus

Trachypithecus_francoisi

Trachypithecus_geei

Trachypithecus_germaini

Trachypithecus_hatinhensis

Trachypithecus_laotum

Trachypithecus_leucocephalus

Trachypithecus_melamera

Trachypithecus_obscurus

Trachypithecus_phayrei

Trachypithecus_pileatus

Reference species: Saguinus_midas

Species mapped:

Leontocebus_fuscicollis

Leontocebus_illigeri

Leontocebus_nigricollis

Saguinus_bicolor

Saguinus_geoffroyi

Saguinus_imperator

Saguinus_inustus

Saguinus_labiatus

Saguinus_midas

Saguinus_mystax

Saguinus_oedipus

Reference species: Sapajus_apella

Species mapped:

Sapajus_apella

Sapajus_macrocephalus

Reference species: Tarsius_syrichta

Species mapped:

Carlito_syrichta

Cephalopachus_bancanus

Tarsius_lariang

Tarsius_wallacei

Reference species: Theropithecus_gelada

Species mapped:

Theropithecus_gelada

LiftOver of the Hg38 annotation to each assembly was performed to obtain GTF files with the coordinates of the coding sequences (CDSs) of each of the genes. The Ensembl ID of the hg38 canonical forms used are in supplementary table S1. The coordinates for each of the references used, and the liftedOver beds are available deposited at Zenodo under the doi 10.5281/zenodo.10637110.

**Supplementary table S1:** Ensembl identifiers of the human canonical isoforms used for this study

| Gene | ENSEMBL ID |
|---|---|
| AHSG | ENST00000411641.7 |
| ALB | ENST00000295897.9 |
| AMBN | ENST00000322937.10 |
| AMELX | ENST00000380714.7 |
| AMTN | ENST00000339336.9 |
| COL17A1 | ENST00000648076.2 |
| COL1A1 | ENST00000225964.10 |
| COL1A2 | ENST00000297268.11 |
| COL2A1 | ENST00000380518.8 |
| ENAM | ENST00000396073.4 |
| MMP20 | ENST00000260228.3 |
| ODAM | ENST00000683306.1 |
| SERPINC1 | ENST00000367698.4 |
| TUFT1 | ENST00000368849.8 |

The VCFs were queried, and the sequences obtained filtered through an in house script using samtools (Li et al. 2009) and bcftools (Li 2011), and then translated using python with standard genetic code.

The genotype calling was performed through the following expression:

```
samtools faidx assembly.fasta ${scaffold}:${start}-${stop} | bcftools consensus –sample ${samplename} -m mask.bed -H 1 file.vcf
```

SNPs were called and filtered using expression:

```
TYPE!='snp' | (GT='het' & FMT/AD[*:*] < 3 ) | AC > 2 | FMT/DP <= 10 | QD < 2 | FS > 60 | INFO/MQ < 40 | SOR > 3 | ReadPosRankSum < -8 | QUAL < 30 | MQRankSum < -12.5
```

And INDELS were called and filtered using:

TYPE!='indel' | (GT='het' & FMT/AD[*:*] < 3 ) | FMT/DP <= 10 | QD < 2 | FS > 200 | INFO/MQ < 40 | SOR > 3 | ReadPosRankSum < -20 | QUAL < 30

At heterozygous positions, bcftools was randomly run either with option –H1 or –H2 for random allele selection.

#### 1.2 Selecting the more comparable sequences for analysis

For the 561 individuals stemming from Kuderna et al. (2023) two sets of translations were performed, one using the original annotation files present in its publication, and the other by using the liftOver coordinates from hg38 reference genome for each of the 32 genomes used (previous section). Each of the sequences for both cases were aligned against the human hg38 canonical form with MAFFT v7.490 (Katoh & Standley, 2013). If only an original, or a liftOver-based translated sequence was available, this sequence was kept for further analysis. If both versions of the sequence were present, gaps of each sequence aligned to the hg38 reference were counted. The sequence that yielded less gappings against the hg38 model was kept for further analysis. The resulting set of sequences is a mix of original and liftOver sequences. A file describing which model was kept for each of the genes per reference is available at Zenodo (10.5281/zenodo.10637110).

#### 1.3 Aligning and refining the sequences

Proteins were aligned using MAFFT v7.490 through the command:

mafft --maxiterate 1000 --globalpair protein.fasta > aligned_protein.fasta

After alignment, a first round of less stringent trimming was performed using the program trimAl 1.2rev59 (Capella-Gutiérrez, Silla-Martínez, and Gabaldón 2009) with the command:

```
trimal -in <inputfile> -out <outputfile> -gt 04 -cons 09
```

For some proteins the value of -gt was lowered from 0.9 to 0.4 and the option -cons was dropped (minimum percentage of the positions in the original alignment to conserve.), resulting in proteins COL17A1 and COL1A1 trimmed with -gt= 0.4, COL2A1 -gt=0.8, ENAM and TUFT1 -gt= 0.9.

After trimming, the proteins were explored through jalview for nonsense variation caused by frameshifts, most likely due to indels. The coordinates of the nonsense variation were manually registered into tsv files available at Zenodo (10.5281/zenodo.10637110). These coordinates were used to find the positions in the DNA sequence causing the frame shift, and edit the DNA in order to recover the frame through an in-house python script.

After recovering the frame of the proteins they were again aligned through the command,

```
mafft --maxiterate 1000 --globalpair protein.fasta > aligned_protein.fasta
```

and trimmed with trimal.

AHSG, ALB, AMBN AMELX, AMTN MMP20, ODAM and SERPINC1 were trimmed with the parameters -gt 09 –cons 60; COL17A1 and COL1A1 with -gt 04; ENAM and TUFT1 with -gt 09.

The resulting trimmed files were further revised through jalview. The regions that still presented spurious variation were manually masked with a “?” symbol. The coordinates of masked regions are available as supplementary file 2.

#### 1.4 Testing the effect of intraspecific variation

In order to not skew the analysis results toward species that are overrepresented with many individuals, most analyses were performed on a multiple sequence alignment (MSA) that contained exactly one individual per species. For each species in this MSA, the individual with the most complete sequences (i.e. least “?”, “X” or “-” characters) was chosen. To test the influence on the outcome, if other, maybe just slightly less complete individuals had been chosen, we created 1000 MSAs by randomly sampling one individual per species (more details in Methods section of the main text). Shannon entropy, Rate4Site score, and a maximum likelihood ML tree (IQ-TREE v. 1.6.12) were calculated for each of the 1000 MSAs and compared to the results generated from the MSA with the most completely sequenced individuals (supplementary figure S1).

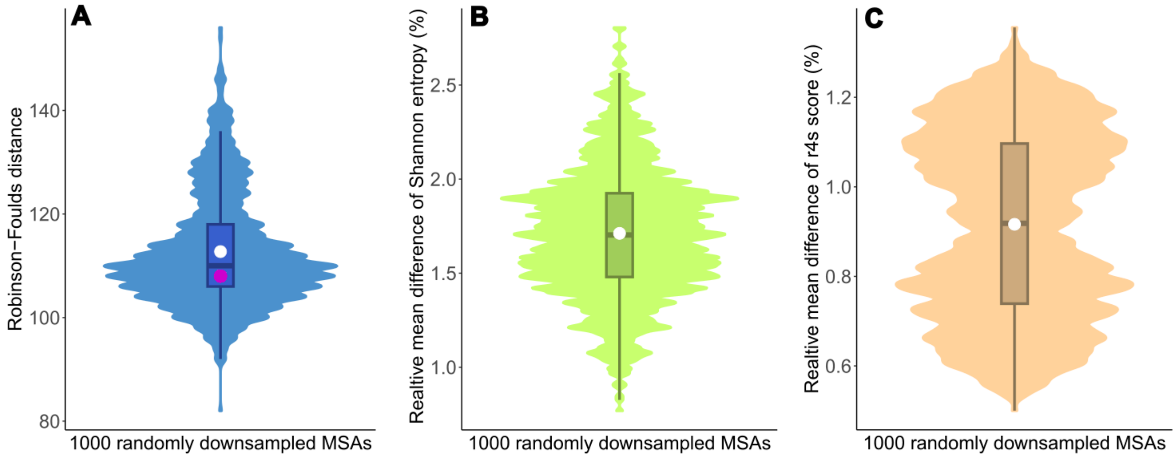

**Supplementary figure S1: Relative Shannon entropy, Rate4Site scores, and Robinson-Foulds distances of 1000 randomly downsamples MSAs** A - Robinson-Foulds distances (RF-distances) to reference species tree (Kuderna et al. 2023), mean is marked as white dot, RF-distance of most complete MSA is marked as pink dot, B - average difference between Shannon entropy at each site of random MSA and most complete MSA as percentage of range of values, C - average difference between Rate4Site score at each site of random MSA and most complete MSA as percentage of range of values

#### 1.4 Reversed phase liquid chromatography tandem mass spectrometry

In order to simulate sequence degradation, we needed to create a simple model of how tooth enamel protein sequences degrade *post mortem*. We mainly used publicly available data (more details see methods in main text), but also added some ancient samples whose processing is described in the following sections.

Samples were analyzed using an Orbitrap Eclipse mass spectrometer (Thermo Fisher Scientific, San Jose, USA) coupled to an EASY-nLC 1200 (Thermo Fisher Scientific, San Jose, USA).

Peptides were loaded directly onto the analytical column and were separated by reversed-phase chromatography using a 50-cm column with an inner diameter of 75 μm, packed with 2 μm C18 particles spectrometer (Thermo Scientific, San Jose, USA). Chromatographic gradients started at 95% buffer A and 5% buffer B with a flow rate of 300 nl/min and gradually increased to 25% buffer B and 75% A in 105 min and then to 40% buffer B and 60% A in 15 min. After each analysis, the column was washed for 10 min with 100% buffer B. Buffer A: 0.1% formic acid in water. Buffer B: 0.1% formic acid in 80% acetonitrile.

The mass spectrometer was operated in positive ionization mode with nanospray voltage set at 2.4 kV and source temperature at 305 °C. The acquisition was performed in data-dependent acquisition (DDA) mode and full MS scans with 1 micro scan at resolution of 120,000 were used over a mass range of m/z 350-1400 with detection in the Orbitrap mass analyzer. Auto gain control (AGC) was set to 4E6 and injection time to 'auto'. In each cycle of data-dependent acquisition analysis, following each survey scan, the most intense ions above a threshold ion count of 10000 were selected for fragmentation. The number of selected precursor ions for fragmentation was determined by the "Top Speed" acquisition algorithm and a dynamic exclusion of 60 seconds. Fragment ion spectra were produced via high-energy collision dissociation (HCD) at normalized collision energy of 28% and they were acquired in the Orbitrap mass analyzer at resolution 30000. AGC was set to 3E4, and an isolation window of 0.7 m/z and a maximum injection time of 54 ms were used.

Four blank samples were injected before and after each sample to avoid sample carryover. Digested bovine serum albumin (New england biolabs cat # P8108S) was analyzed between each sample to avoid sample carryover and to assure stability of the instrument and QCloud (Chiva et al. 2018) has been used to control instrument longitudinal performance during the project.

#### 1.5 Ancient protein sequence reconstruction

Equid samples were run using MaxQuant v.1.6.17.0, with the following settings different from default as described in supplementary table S2. The reference equid enamelome database was mainly built from publicly available protein sequences of ALB, COL1A2, COL1A1, COL2A1, COL17A1, AMBN, AMELX, ENAM, AMELY, AMTN, MMP20, KLK4, TUFT1, SERPINC1 and ODAM from UniProt and GenBank. All isoforms were included.

Deinotheriid samples were run using MaxQuant v2.0.2.0, including MaxNovo, with the following settings different from default as described in supplementary table S2.

The reference Proboscidea enamelome database was built using ProteoParc v1.0. The TaxID was set as 9779 (Order Proboscidea) and the search was focused on the above described teeth proteins: ALB, COL1A2, COL1A1, COL2A1, COL17A1, AMBN, AMELX, ENAM, AMELY, AMTN, MMP20, KLK4, TUFT1, SERPINC1 and ODAM.

Ancient sequence reconstruction was performed grouping the peptide sequences by the most common “Leading razor protein” in each protein group. Reverse and contaminant peptides were removed from the final reconstruction during this step.

**Supplementary table S2:** MaxQuant parameter settings. Carbamidomethyl (C) was removed from the fixed modifications parameter as the sample preparation does not produce this type of chemical conversion.

| **Parameter** | **Setting v.1.6.17.0** | **Setting v.2.0.2.0** |
|---|---|---|
| Digestion | Unspecific | Unspecific |
| Fixed modifications | None | None |
| Variable modifications | Oxidation of M; Oxidation of P; Gln->Pyro-Glu; Glu->Pyro-Glu; Deamidation (NQ); Phosphorylation (ST) | Oxidation of M; Oxidation of P; Gln->Pyro-Glu; Glu->Pyro-Glu; Deamidation (NQ); Phosphorylation (ST) |
| Maximum number of modifications per peptide | 5 | 4 |
| Main search peptide tolerance | 4.5 ppm | 10 ppm |
| Individual peptide mass tolerance | TRUE | FALSE |
| Minimum peptide length for unspecific search | 7 | 7 |
| Protein FDR | 1 | 1 |
| Dependent peptides | FALSE | TRUE |
| De-novo sequencing | FALSE | TRUE |
| Minimum delta score for modified peptides | 0 | 0 |
| Minimum score for unmodified peptides | 40 | 40 |
| use .NET Core | TRUE | FALSE (for windows OS) |

### 2. Concatenations and Phylogenetic analysis

The signal peptide was removed, by trimming the following number of peptides from the beginning of the sequence of each protein, following the signal peptide sequence reported at UniProt:

| | |
|---|---|
| AHSG | 17 |
| ALB | 17 |
| AMBN | 21 |
| AMTN | 16 |
| COL17A1 | 0 |
| COL1A1 | 22 |
| COL1A2 | 22 |
| COL2A1 | 25 |
| ENAM | 39 |
| ODAM | 15 |
| MMP20 | 22 |
| SERPINC1 | 32 |
| TUFT1 | 20 |

The trimmed sequences were concatenated into three different groups of 5, 10 and 14 proteins with the command:

```
$ catfasta2phyml.pl prot1.fasta prot2.fasta (...) protX.fasta > concatenation.fasta
```

The three concatenations that were performed are:

5 proteins: AMBN, AMELX, AMTN, ENAM, MMP20

10 proteins: AHSG, ALB, AMBN, AMELX, AMTN, ENAM, MMP20, ENAM, ODAM, SERPINC1

14 proteins: AHSG, ALB, AMBN, AMELX, AMTN, COL17A1, COL1A1, COL1A2, COL2A1, ENAM, MMP20, ODAM, SERPINC1

ML and Bayesian phylogenetic analyses were performed for the three concatenations.

For the phylogenetic analysis, the dataset was reduced to one individual per species, yielding a dataset of 233 individuals.

The evolutionary model of each of the proteins was obtained using model finder (Kalyaanamoorthy et al. 2017) with the command:

```
iqtree -s Aligned_protein_file.fasta -nt 6 -redo -m MF -msub nuclear
```

yielding the following models:

AHSG = JTT+R3

ALB = JTTDCMut+G4

AMBN = JTT+I+G4

AMELX = JTTDCMut+F+G4

AMTN = JTT+R2

COL17A1 = JTT+R3

COL1A1 = JTTDCMut+F+R3

COL1A2 = JTTDCMut+F+R3

COL2A1 = JTTDCMut+F+R3

ENAM = JTT+R3

MMP20 = JTT+I+G4

ODAM = JTT+I+G4

SERPINC1 = JTT+I+G4

TUFT1 = JTT+G4

ML phylogenetic analysis was performed with IQ-TREE1.6.12 (Nguyen et al. 2015) through the command:

```
iqtree -nt 4 -s $infile -spp $partfile -alrt 5000 -bb 5000 -nstop 500 -nm 10000 -wsplits -pre $outdir
```

Bayesian analysis was performed with MrBayes v.3.2.7a (Ronquist and Huelsenbeck 2003) for 3 million generations with the parameters:

```
rates=invgamma nucmodel=Protein;
unlink statefreq=(all) revmat=(all) shape=(all) pinvar=(all);
prset applyto=(all) ratepr=variable aamodel=mixed;
temp=0.2
Ngen=3000000
Diagnfreq=5000
Nruns=2
Nchains=4
relburnin=yes
Burninfrac=0.25
samplefreq=500
printfreq=500
starttree=random
savebrlens=yes
startparams=reset xx x
Filename=out_files/14_proteins/14_proteins.out;
```

Convergence was assessed with TRACER (Rambaut et al. 2018) (output available at 10.5281/zenodo.10637110)

The ML trees based on the 5 and 10 protein concatenations are displayed in supplementary figure S2.

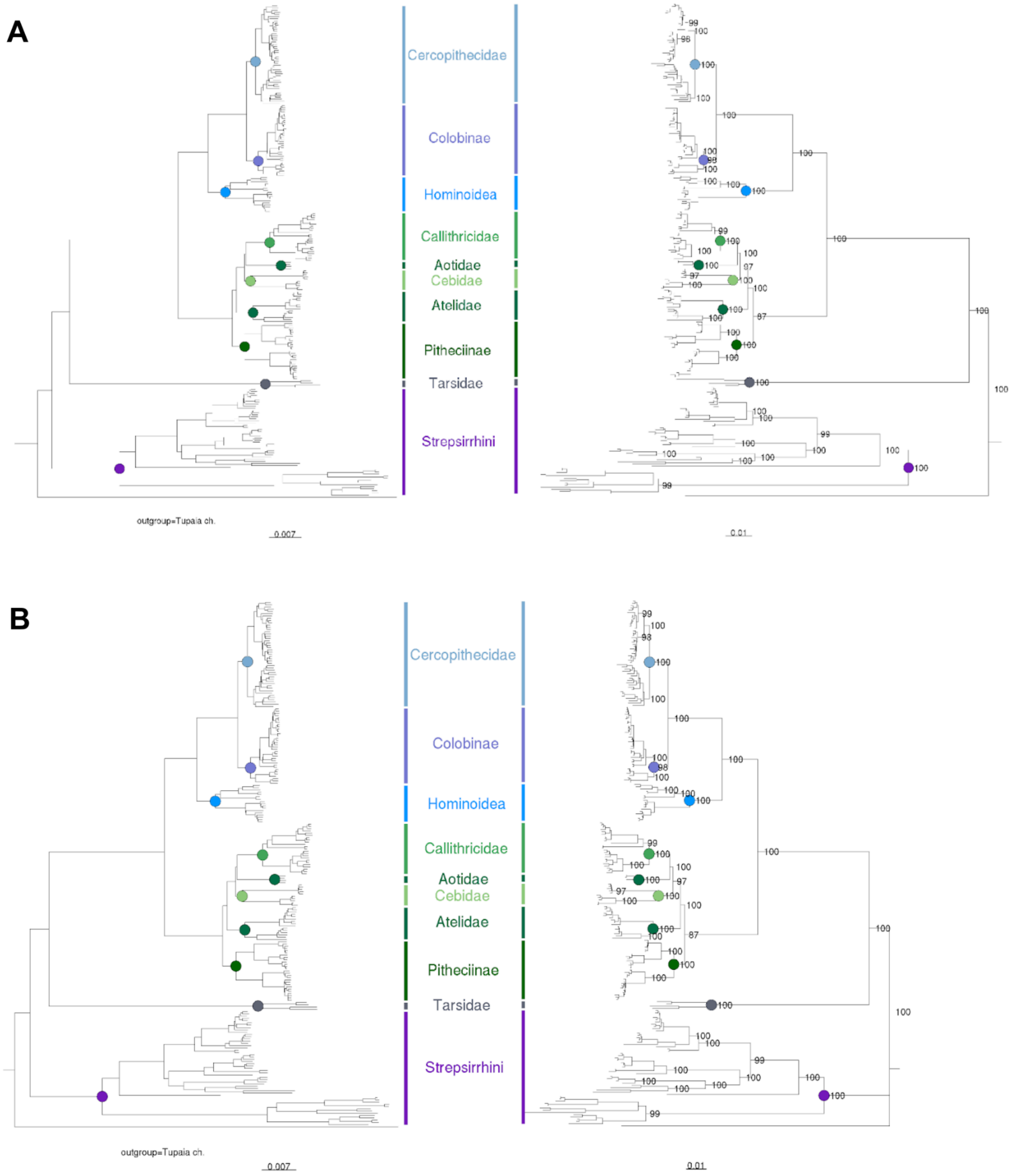

**Supplementary figure S2: Phylogenies based on concatenation of five and ten proteins**

Created using ML (IQ-TREE v1.6.12). A - five proteins, B - ten proteins. Trees on the left are the species tree, trees on the right the phylogenies computed with the concatenations.

In the 14 protein concatenation-based phylogenies, which contain collagens, it was seen that the tarsiers appeared wrongly positioned as monophyletic with strepsirrhines (see Results in main text). For this reason, we performed phylogenetic analysis on each of the collagens and in all the permutations possible of collagens along with the complete set of non collagen proteins (10 proteins). This approach allowed us to explore which of the proteins were driving the observed placement of the tarsiers in the trees based on the 14 protein concatenations. We can see in supplementary table S3 in which sequences or combinations of sequences the tarsiers appear correctly (i.e. "sister to simiiformes") or incorrectly positioned.

**Supplementary table S3:** Phylogenetic position of tarsiers when computing ML with the selection of protein sequences displayed in column 1. "E" stands for enamel and represents 10 non-collagenous enamel proteins (AHSG, ALB, AMBN, AMELX, AMTN ENAM, MMP20, ODAM, SERPINC1, TUFT1).

| **Enamel and Collagens in dataset** | **Positioning of Tarsiers** | **Bootstrap value** |
|---|---|---|
| COL17A1 | sister to strepsirrhines | 55 |
| COL1A1 | sister to strepsirrhines with lorises and galagos | 92 |
| COL1A2 | sister to strepsirrhines | 98 |
| COL2A1 | sister to simiiformes, with strepsirrhines unresolved | 100 |
| 4 collagens | sister to strepsirrhines | 99 |
| Enamel (E) + 4 collagens | sister to strepsirrhines | 90 |
| E + COL17A1, COL1A2 | sister to strepsirrhines | 73 |
| E + COL1A1, COL1A2 | sister to strepsirrhines | 78 |
| E + COL17A1, COL1A1, COL1A2 | sister to strepsirrhines | 85 |
| E + COL1A1, COL2A1, COL1A2 | sister to strepsirrhines | 78 |
| E + COL17A1, COL1A2, COL2A1 | sister to strepsirrhines | 72 |
| E + COL17A1, COL1A1 | sister to simiiformes | 100 |
| E + COL17A1, COL2A1 | sister to simiiformes | 95 |
| E + COL1A1, COL2A1 | sister to simiiformes | 95 |
| E + COL1A2, COL2A1 | sister to simiiformes | 93 |
| E + COL17A1, COL1A1, COL2A1 | sister to simiiformes | 88 |
| E + COL17A1 | sister to simiiformes | 95 |
| E + COL1A1 | sister to simiiformes | 98 |
| E + COL1A2 | sister to simiiformes | 93 |
| E + COL2A1 | sister to simiiformes | 100 |

### 3. Rate4Site scores and Shannon entropy

The degree of sequence conservation of each of the amino acid sites through the five, ten and 14 protein concatenations was estimated in two ways. First, with Shannon entropy with the script Shannon.py (https://gist.github.com/jrjhealey). Given the fact that this metric is agnostic to evolutionary constraints, Rate4Site scores (Pupko et al. 2002) were also calculated. Rate4Site first computes a phylogenetic tree and calculates the rate of each site considering the topology and the length of the branches. Rate4Site was applied through the debian package Rate4Site (https://debian.pkgs.org/10/debian-main-arm64/Rate4Site_3.0.0-6_arm64.deb.html).

For Shannon entropy, the computation was performed on the fasta files of concatenated MSAs. In contrast, for Rate4Site, the computation was made on both, single proteins (used to divide into variable and conserved datasets) and the concatenated MSA (used for all other figures). Both metrics along the three protein concatenations (5, 10, and 14 proteins) were then normalized to mean of 1. The output files are available at Zenodo (10.5281/zenodo.10637110).

Based on the Rate4Site and Shannon entropy scores we built files for each of the three concatenations (5, 10 and 14 proteins) by subsetting them by values over and under the threshold of 1. The files with values over this threshold were tagged as “variable”, and the ones with the values under the threshold, as “conserved” (supplementary figure S3).

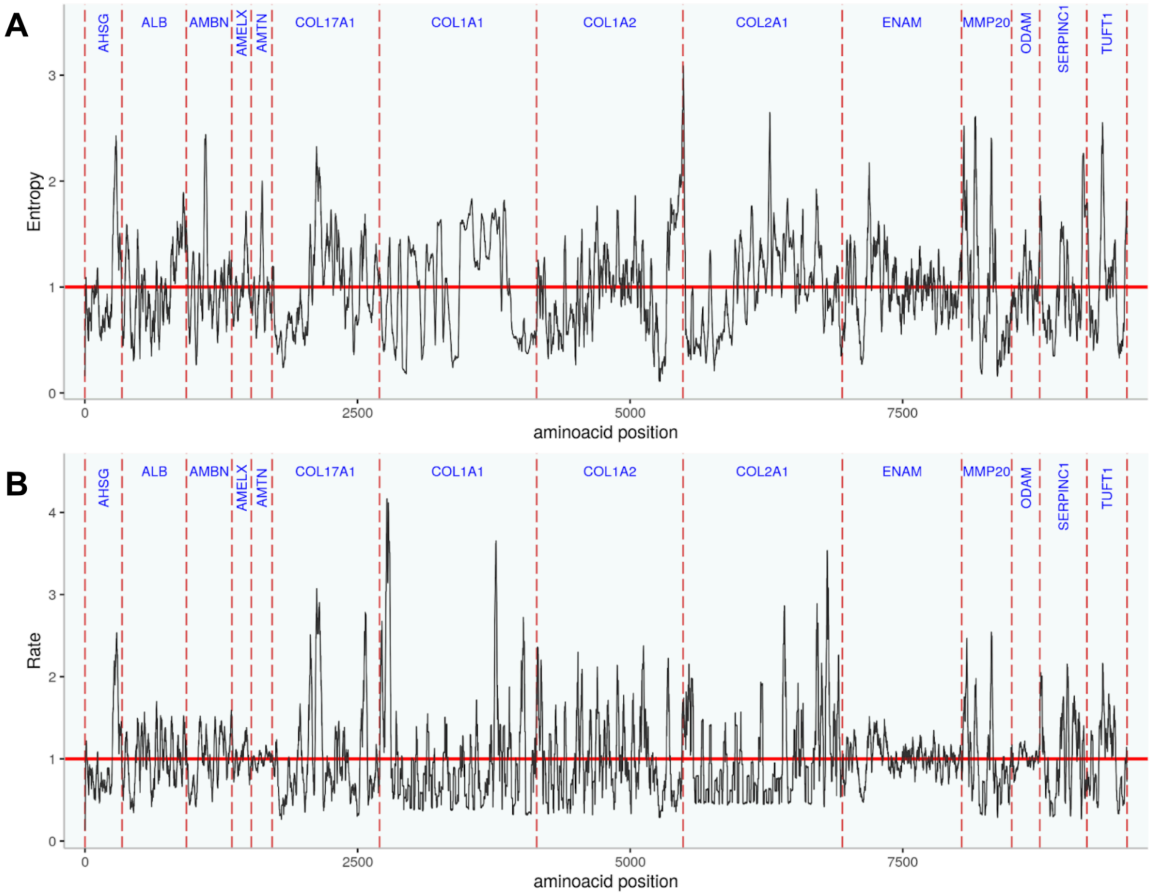

**Supplementary figure S3: Division of each protein into “conserved” and “variable” sites**

Sites with Shannon entropy or Rate4Site values below 1 are considered “conserved”, sites above this value “variable”. The calculation of these values and normalization has been performed on each protein individually before concatenating the “variable” and “conserved” sites into an MSA, respectively and for each method and concatenation (5,10, and 14 proteins), which resulted in twelve MSAs. A - by Shannon entropy; B - by Rate4Site score

The result was twelve fasta files for each scoring system, two for each variability state. The subsetting was performed through in-house python code. We computed ML phylogenetic analysis over each file with IQ-TREE v1.6.12 through the command:

```
iqtree -nt 4 -s $infile -spp $partfile -alrt 5000 -bb 5000 -nstop 500 -nm 10000 -wsplits -pre $outdir
```

Where the amino acid substitution models used for the analysis for each protein partition was the same as reported in the section "Concatenations and Phylogenetic Analysis".

We calculated RF-distances for each of the twelve files against the species tree reported in Kuderna et al. (2023) through the function treedist of the R package phangorn (https://cran.r-project.org/web/packages/phangorn/index.html).

To compare the patterns of sequence conservation of primates, Rate4Site scores were calculated for a set of mammals. The species were chosen in a manner to represent each branch of the mammalian lineage. Species and exact sequences used for the calculations are noted in supplementary table S4. The resulting Rate4Site scores are displayed in supplementary figure S4.

**Supplementary table S4:** Protein identifiers and species of mammals used to calculate Rate4Site scores. In AMBN and AMELX, a hybrid of *Bos taurus x Bos indicus* was used instead of *Bos taurus* because of better sequence quality.

| UniParc ID | Genus | Species Epitheton | Gene |
|---|---|---|---|
| UPI00025DC9FE | *Ictidomys* | *tridecemlineatus* | AHSG |
| UPI0001F16D78 | *Ailuropoda* | *melanoleuca* | AHSG |
| UPI0003EE3D23 | *Leptonychotes* | *weddellii* | AHSG |
| UPI00029907A1 | *Felis* | *catus* | AHSG |
| UPI001E26DF69 | *Lemur* | *catta* | AHSG |
| UPI000D184883 | *Desmodus* | *rotundus* | AHSG |
| UPI0001C5B6FA | *Loxodonta* | *africana* | AHSG |
| UPI00045E4D8F | *Orycteropus* | *afer* | AHSG |
| UPI0003AFC8EB | *Capra* | *hircus* | AHSG |
| UPI0023FAA5B1 | *Ovis* | *aries* | AHSG |
| UPI0000125015 | *Bos* | *taurus* | AHSG |
| UPI001346041F | *Moschus* | *moschiferus* | AHSG |
| UPI000BB8DBA5 | *Delphinapterus* | *leucas* | AHSG |
| UPI0003C88453 | *Camelus* | *ferus* | AHSG |
| UPI000333C952 | *Echinops* | *telfairi* | AHSG |
| UPI0000125017 | *Homo* | *sapiens* | AHSG |
| UPI0001FB1964 | *Equus* | *caballus* | AHSG |
| UPI000000191F | *Mus* | *musculus* | AHSG |

| | | | |
|---|---|---|---|
| UPI00038BD484 | *Microtus* | *ochrogaster* | AHSG |
| UPI000328F00F | *Dasypus* | *novemcinctus* | AHSG |
| UPI0000F63D7F | *Erinaceus* | *europaeus* | AHSG |
| UPI0003343EAD | *Condylura* | *cristata* | AHSG |
| UPI00001257C2 | *Felis* | *catus* | ALB |
| UPI0001DECA0B | *Ailuropoda* | *melanoleuca* | ALB |
| UPI0003EE3118 | *Leptonychotes* | *weddellii* | ALB |
| UPI000D186FF9 | *Desmodus* | *rotundus* | ALB |
| UPI0003346446 | *Condylura* | *cristata* | ALB |
| UPI000004ECF8 | *Bos* | *taurus* | ALB |
| UPI00001257CB | *Ovis* | *aries* | ALB |
| UPI000BB6C4F8 | *Delphinapterus* | *leucas* | ALB |
| UPI0003C8A000 | *Camelus* | *ferus* | ALB |
| UPI000002C1AC | *Homo* | *sapiens* | ALB |
| UPI001C5C884A | *Orycteropus* | *afer* | ALB |
| UPI001E26D6CF | *Lemur* | *catta* | ALB |
| UPI00001257C3 | *Equus* | *caballus* | ALB |
| UPI0001C5F149 | *Loxodonta* | *africana* | ALB |
| UPI000443926B | *Erinaceus* | *europaeus* | ALB |
| UPI00000011E3 | *Mus* | *musculus* | ALB |
| UPI00038C3C0B | *Microtus* | *ochrogaster* | ALB |
| UPI0003CC0A0C | *Dasypus* | *novemcinctus* | ALB |
| UPI00025DA330 | *Ictidomys* | *tridecemlineatus* | ALB |
| consensus_UPI001343547A_UPI001334EB93 | *Moschus* | *moschiferus* | ALB |
| consensus_UPI0000E43A39_UPI001E1E0F8E | *Echinops* | *telfairi* | ALB |
| UPI0003AF5A5F | *Capra* | *hircus* | ALB |
| UPI00025DA2E2 | *Ictidomys* | *tridecemlineatus* | AMBN |
| UPI000C812A7E | *Loxodonta* | *africana* | AMBN |
| UPI001E267C92 | *Lemur* | *catta* | AMBN |
| UPI00045DB8A8 | *Orycteropus* | *afer* | AMBN |
| UPI00004A0469 | *Mus* | *musculus* | AMBN |
| UPI0003EDC605 | *Leptonychotes* | *weddellii* | AMBN |
| UPI0002AD4D39 | *Felis* | *catus* | AMBN |
| UPI000BB71DA7 | *Delphinapterus* | *leucas* | AMBN |
| UPI00038BF908 | *Microtus* | *ochrogaster* | AMBN |
| UPI0013A62954 | *Camelus* | *ferus* | AMBN |
| UPI001235A039 | *Equus* | *caballus* | AMBN |
| UPI00029D6FC5 | *Ovis* | *aries* | AMBN |
| UPI00059B09E9 | *Ailuropoda* | *melanoleuca* | AMBN |

| | | | |
|---|---|---|---|
| UPI0003AFBAE5 | *Capra* | *hircus* | AMBN |
| UPI0003343AF3 | *Condylura* | *cristata* | AMBN |
| UPI0006D8F6B5 | *Bos* | *taurus x indicus* | AMBN |
| UPI00133DC85A | *Moschus* | *moschiferus* | AMBN |
| UPI000000DCCB | *Homo* | *sapiens* | AMBN |
| UPI000333E489 | *Echinops* | *telfairi* | AMBN |
| UPI000D184425 | *Desmodus* | *rotundus* | AMBN |
| UPI000328CF7A | *Dasypus* | *novemcinctus* | AMBN |
| UPI000444297F | *Erinaceus* | *europaeus* | AMBN |
| UPI00000E705F | *Mus* | *musculus* | AMELX |
| UPI00038C31DD | *Microtus* | *ochrogaster* | AMELX |
| UPI0001510A3B | *Ictidomys* | *tridecemlineatus* | AMELX |
| UPI0000E41C37 | *Echinops* | *telfairi* | AMELX |
| UPI0001C5D780 | *Loxodonta* | *africana* | AMELX |
| UPI000002A3B7 | *Homo* | *sapiens* | AMELX |
| UPI0000F6350E | *Erinaceus* | *europaeus* | AMELX |
| UPI000D1810BA | *Desmodus* | *rotundus* | AMELX |
| UPI000057EE88 | *Bos* | *taurus x indicus* | AMELX |
| UPI0006B12AE5 | *Capra* | *hircus* | AMELX |
| UPI00134C2B05 | *Moschus* | *moschiferus* | AMELX |
| UPI00194C6239 | *Ailuropoda* | *melanoleuca* | AMELX |
| UPI000DE7D14B | *Felis* | *catus* | AMELX |
| UPI00057B080A | *Camelus* | *ferus* | AMELX |
| UPI00033446DD | *Condylura* | *cristata* | AMELX |
| UPI001235B980 | *Equus* | *caballus* | AMELX |
| UPI0003EDE479 | *Leptonychotes* | *weddellii* | AMELX |
| UPI001E26D384 | *Lemur* | *catta* | AMELX |
| UPI0000DB81B5 | *Ovis* | *aries* | AMELX |
| UPI00062AA047 | *Dasypus* | *novemcinctus* | AMELX |
| UPI00122CF1E6 | *Delphinapterus* | *leucas* | AMELX |
| UPI00045DD7A1 | *Orycteropus* | *afer* | AMELX |
| H2PUX0 | *Pongo* | *abelii* | AMELX |
| F6PLF2 | *Canis* | *lupus* | AMELX |
| A0A1D5RDA1 | *Macaca* | *mulatta* | AMELY |
| Q99218 | *Homo* | *sapiens* | AMELY |
| Q861X8 | *Pan* | *troglodytes* | AMELY |
| W8CEL7 | *Gorilla* | *gorilla* | AMELY |
| Q99004 | *Bos* | *taurus* | AMELY |
| Q9TUI2 | *Equus* | *caballus* | AMELY |
| Q861X6 | *Saimiri* | *sciureus* | AMELY |
| Q861W9 | *Sus* | *scrofa* | AMELY |
| UPI00064E6D0F | *Echinops* | *telfairi* | AMTN |

| | | | |
|---|---|---|---|
| UPI000BB67AC4 | *Delphinapterus* | *leucas* | AMTN |
| UPI00045DAB7C | *Orycteropus* | *afer* | AMTN |
| UPI0001C5F2F4 | *Loxodonta* | *africana* | AMTN |
| UPI00059AE37A | *Ailuropoda* | *melanoleuca* | AMTN |
| UPI00057B3196 | *Camelus* | *ferus* | AMTN |
| UPI0003ACA174 | *Equus* | *caballus* | AMTN |
| UPI00000389F3 | *Homo* | *sapiens* | AMTN |
| UPI0003344074 | *Condylura* | *cristata* | AMTN |
| UPI00134501CB | *Moschus* | *moschiferus* | AMTN |
| UPI001C2E0034 | *Ovis* | *aries* | AMTN |
| UPI0003F1B638 | *Felis* | *catus* | AMTN |
| UPI0000EBF097 | *Bos* | *taurus* | AMTN |
| UPI000BADC7EA | *Ictidomys* | *tridecemlineatus* | AMTN |
| UPI0000029DDC | *Mus* | *musculus* | AMTN |
| UPI0003AFA8CF | *Capra* | *hircus* | AMTN |
| UPI00238146D9 | *Desmodus* | *rotundus* | AMTN |
| UPI0003EDF5CC | *Leptonychotes* | *weddellii* | AMTN |
| UPI001E26A062 | *Lemur* | *catta* | AMTN |
| UPI00038C4533 | *Microtus* | *ochrogaster* | AMTN |
| UPI000C837699 | *Dasypus* | *novemcinctus* | AMTN |
| UPI0000F61C6D | *Erinaceus* | *europaeus* | AMTN |
| UPI0014946839 | *Ailuropoda* | *melanoleuca* | COL17A1 |
| UPI00015885B6 | *Bos* | *taurus* | COL17A1 |
| UPI0013A681D7 | *Camelus* | *ferus* | COL17A1 |
| UPI000846E3F9 | *Capra* | *hircus* | COL17A1 |
| UPI000643CF4E | *Condylura* | *cristata* | COL17A1 |
| UPI00062A778E | *Dasypus* | *novemcinctus* | COL17A1 |
| UPI000BB74BB4 | *Delphinapterus* | *leucas* | COL17A1 |
| UPI000D1822A7 | *Desmodus* | *rotundus* | COL17A1 |
| UPI00064EC0AD | *Echinops* | *telfairi* | COL17A1 |
| UPI000C9E8A63 | *Equus* | *caballus* | COL17A1 |
| UPI000443BA49 | *Erinaceus* | *europaeus* | COL17A1 |
| UPI001D19C19E | *Felis* | *catus* | COL17A1 |
| UPI000006DB58 | *Homo* | *sapiens* | COL17A1 |
| UPI00068077E8 | *Ictidomys* | *tridecemlineatus* | COL17A1 |
| UPI001E26B01E | *Lemur* | *catta* | COL17A1 |
| UPI0003EDDE6C | *Leptonychotes* | *weddellii* | COL17A1 |
| UPI000C813626 | *Loxodonta* | *africana* | COL17A1 |
| UPI00067CEC6C | *Microtus* | *ochrogaster* | COL17A1 |
| UPI00134886E1 | *Moschus* | *moschiferus* | COL17A1 |
| UPI0000475485 | *Mus* | *musculus* | COL17A1 |
| UPI001C5CB292 | *Orycteropus* | *afer* | COL17A1 |

| | | | |
|---|---|---|---|
| UPI0005FAA81D | *Ovis* | *aries* | COL17A1 |
| UPI000C9EBA1E | *Equus* | *caballus* | COL1A1 |
| UPI000003BB1E | *Bos* | *taurus* | COL1A1 |
| UPI0003EE2841 | *Leptonychotes* | *weddellii* | COL1A1 |
| UPI0000020B83 | *Mus* | *musculus* | COL1A1 |
| UPI0001DEBE30 | *Ailuropoda* | *melanoleuca* | COL1A1 |
| UPI000298B65B | *Felis* | *catus* | COL1A1 |
| UPI001345F54E | *Moschus* | *moschiferus* | COL1A1 |
| UPI0005FBADFE | *Ovis* | *aries* | COL1A1 |
| UPI000847268D | *Capra* | *hircus* | COL1A1 |
| UPI0000DACAC3 | *Homo* | *sapiens* | COL1A1 |
| UPI00070453D3 | *Camelus* | *ferus* | COL1A1 |
| UPI0005406912 | *Loxodonta* | *africana* | COL1A1 |
| UPI001A9DA007 | *Ictidomys* | *tridecemlineatus* | COL1A1 |
| UPI000BB84A00 | *Delphinapterus* | *leucas* | COL1A1 |
| UPI0003336B52 | *Echinops* | *telfairi* | COL1A1 |
| UPI001E2667D5 | *Lemur* | *catta* | COL1A1 |
| UPI000334613F | *Condylura* | *cristata* | COL1A1 |
| UPI0023812E63 | *Desmodus* | *rotundus* | COL1A1 |
| UPI00045E115D | *Orycteropus* | *afer* | COL1A1 |
| UPI0004439EB6 | *Erinaceus* | *europaeus* | COL1A1 |
| UPI000328F48D | *Dasypus* | *novemcinctus* | COL1A1 |
| UPI00038BEBF8 | *Microtus* | *ochrogaster* | COL1A1 |
| UPI0001DEAD0B | *Ailuropoda* | *melanoleuca* | COL1A2 |
| UPI0000126D35 | *Bos* | *taurus* | COL1A2 |
| UPI0003C82C5C | *Camelus* | *ferus* | COL1A2 |
| UPI0003AF943D | *Capra* | *hircus* | COL1A2 |
| UPI00033455AE | *Condylura* | *cristata* | COL1A2 |
| UPI0003292562 | *Dasypus* | *novemcinctus* | COL1A2 |
| UPI000BB97DA2 | *Delphinapterus* | *leucas* | COL1A2 |
| UPI000D181AB4 | *Desmodus* | *rotundus* | COL1A2 |
| UPI000333F1DE | *Echinops* | *telfairi* | COL1A2 |
| UPI0001FB3191 | *Equus* | *caballus* | COL1A2 |
| UPI000443F7C7 | *Erinaceus* | *europaeus* | COL1A2 |
| UPI000298A177 | *Felis* | *catus* | COL1A2 |
| UPI00001B0786 | *Homo* | *sapiens* | COL1A2 |
| UPI00038C5333 | *Ictidomys* | *tridecemlineatus* | COL1A2 |
| UPI001E26842F | *Lemur* | *catta* | COL1A2 |
| UPI0003EDCB05 | *Leptonychotes* | *weddellii* | COL1A2 |
| UPI0001C5C769 | *Loxodonta* | *africana* | COL1A2 |
| UPI00038C1845 | *Microtus* | *ochrogaster* | COL1A2 |
| UPI001346D826 | *Moschus* | *moschiferus* | COL1A2 |

| | | | |
|---|---|---|---|
| UPI0000044DC6 | *Mus* | *musculus* | COL1A2 |
| UPI00045E253C | *Orycteropus* | *afer* | COL1A2 |
| UPI00029D4E7D | *Ovis* | *aries* | COL1A2 |
| UPI0001F1A265 | *Ailuropoda* | *melanoleuca* | COL2A1 |
| UPI000162E87B | *Bos* | *taurus* | COL2A1 |
| UPI0013A67CAE | *Camelus* | *ferus* | COL2A1 |
| UPI000846A11F | *Capra* | *hircus* | COL2A1 |
| UPI0003346F70 | *Condylura* | *cristata* | COL2A1 |
| UPI000328E64E | *Dasypus* | *novemcinctus* | COL2A1 |
| UPI000BB74C37 | *Delphinapterus* | *leucas* | COL2A1 |
| UPI000D183C63 | *Desmodus* | *rotundus* | COL2A1 |
| UPI000333750D | *Echinops* | *telfairi* | COL2A1 |
| UPI0001FB2989 | *Equus* | *caballus* | COL2A1 |
| UPI0004439497 | *Erinaceus* | *europaeus* | COL2A1 |
| UPI00029890C4 | *Felis* | *catus* | COL2A1 |
| UPI0000D79713 | *Homo* | *sapiens* | COL2A1 |
| UPI00038C63F9 | *Ictidomys* | *tridecemlineatus* | COL2A1 |
| UPI001E266799 | *Lemur* | *catta* | COL2A1 |
| UPI0003EE2F09 | *Leptonychotes* | *weddellii* | COL2A1 |
| UPI0002233784 | *Loxodonta* | *africana* | COL2A1 |
| UPI00038BE84A | *Microtus* | *ochrogaster* | COL2A1 |
| UPI0013431802 | *Moschus* | *moschiferus* | COL2A1 |
| UPI000043A9B1 | *Mus* | *musculus* | COL2A1 |
| UPI00045E51C7 | *Orycteropus* | *afer* | COL2A1 |
| UPI001C2E8633 | *Ovis* | *aries* | COL2A1 |
| UPI0004439DC5 | *Erinaceus* | *europaeus* | ENAM |
| UPI0001DEB032 | *Ailuropoda* | *melanoleuca* | ENAM |
| UPI0003EDE520 | *Leptonychotes* | *weddellii* | ENAM |
| UPI000298CC11 | *Felis* | *catus* | ENAM |
| UPI000D184420 | *Desmodus* | *rotundus* | ENAM |
| UPI000D195468 | *Capra* | *hircus* | ENAM |
| UPI0003CD1C6A | *Ovis* | *aries* | ENAM |
| UPI000FC4D3F9 | *Bos* | *taurus* | ENAM |
| UPI001350ABB9 | *Moschus* | *moschiferus* | ENAM |
| A0A8B8R9K1 | *Camelus* | *ferus* | ENAM |
| UPI0023FBDC02 | *Delphinapterus* | *leucas* | ENAM |
| UPI001E26D3EE | *Lemur* | *catta* | ENAM |
| UPI000013CE60 | *Homo* | *sapiens* | ENAM |
| UPI000155E1FB | *Equus* | *caballus* | ENAM |
| UPI0003343607 | *Condylura* | *cristata* | ENAM |
| UPI0001C5F2ED | *Loxodonta* | *africana* | ENAM |
| UPI00045D6081 | *Orycteropus* | *afer* | ENAM |

| | | | |
|---|---|---|---|
| UPI00025DA2DB | *Ictidomys* | *tridecemlineatus* | ENAM |
| UPI000333824D | *Echinops* | *telfairi* | ENAM |
| UPI000328B984 | *Dasypus* | *novemcinctus* | ENAM |
| UPI000154CC5B | *Mus* | *musculus* | ENAM |
| UPI00038BB978 | *Microtus* | *ochrogaster* | ENAM |
| UPI0000000896 | *Homo* | *sapiens* | KLK4 |
| UPI001E26E4EA | *Lemur* | *catta* | KLK4 |
| UPI001493E8BC | *Ailuropoda* | *melanoleuca* | KLK4 |
| UPI0003EDF9BD | *Leptonychotes* | *weddellii* | KLK4 |
| UPI001D19BEAC | *Felis* | *catus* | KLK4 |
| UPI000846C44C | *Capra* | *hircus* | KLK4 |
| UPI0005FB8487 | *Ovis* | *aries* | KLK4 |
| UPI001333A6C8 | *Moschus* | *moschiferus* | KLK4 |
| UPI00017C3809 | *Bos* | *taurus* | KLK4 |
| UPI001C2E209E | *Ovis* | *aries* | KLK4 |
| UPI0001FB1F6D | *Equus* | *caballus* | KLK4 |
| UPI001A9F0922 | *Ictidomys* | *tridecemlineatus* | KLK4 |
| UPI0004439D51 | *Erinaceus* | *europaeus* | KLK4 |
| UPI0000020FE9 | *Mus* | *musculus* | KLK4 |
| UPI00038C2237 | *Microtus* | *ochrogaster* | KLK4 |
| UPI0003347776 | *Condylura* | *cristata* | KLK4 |
| UPI00038C593F | *Ictidomys* | *tridecemlineatus* | MMP20 |
| UPI001E26A080 | *Lemur* | *catta* | MMP20 |
| UPI0003AF98CD | *Capra* | *hircus* | MMP20 |
| UPI0005FB8FE4 | *Ovis* | *aries* | MMP20 |
| UPI001348900A | *Moschus* | *moschiferus* | MMP20 |
| UPI000012F256 | *Bos* | *taurus* | MMP20 |
| UPI0003C89840 | *Camelus* | *ferus* | MMP20 |
| UPI000BB801D3 | *Delphinapterus* | *leucas* | MMP20 |
| UPI000155E9D4 | *Equus* | *caballus* | MMP20 |
| UPI0004444AF6 | *Erinaceus* | *europaeus* | MMP20 |
| UPI0003344CA9 | *Condylura* | *cristata* | MMP20 |
| UPI001D19FDC5 | *Felis* | *catus* | MMP20 |
| UPI0001C5B5D5 | *Loxodonta* | *africana* | MMP20 |
| UPI000D186A4F | *Desmodus* | *rotundus* | MMP20 |
| UPI00033371A4 | *Echinops* | *telfairi* | MMP20 |
| UPI0014944DE9 | *Ailuropoda* | *melanoleuca* | MMP20 |
| UPI00123EADAD | *Leptonychotes* | *weddellii* | MMP20 |
| UPI00038BD861 | *Microtus* | *ochrogaster* | MMP20 |
| UPI0000023950 | *Mus* | *musculus* | MMP20 |
| UPI00019549AB | *Dasypus* | *novemcinctus* | MMP20 |
| UPI00045D67AA | *Orycteropus* | *afer* | MMP20 |

| | | | |
|---|---|---|---|
| UPI000013D0B3 | *Homo* | *sapiens* | MMP20 |
| UPI0014947B96 | *Ailuropoda* | *melanoleuca* | ODAM |
| UPI0000EAFE61 | *Bos* | *taurus* | ODAM |
| UPI0013A666AA | *Camelus* | *ferus* | ODAM |
| UPI0003AFF1F9 | *Capra* | *hircus* | ODAM |
| UPI0003343C2B | *Condylura* | *cristata* | ODAM |
| UPI0003CC0FFF | *Dasypus* | *novemcinctus* | ODAM |
| UPI000BB7081C | *Delphinapterus* | *leucas* | ODAM |
| UPI001E1C196E | *Desmodus* | *rotundus* | ODAM |
| UPI0003336874 | *Echinops* | *telfairi* | ODAM |
| UPI0001560BCD | *Equus* | *caballus* | ODAM |
| UPI000443CB73 | *Erinaceus* | *europaeus* | ODAM |
| UPI0009484FF4 | *Felis* | *catus* | ODAM |
| UPI000387D1B9 | *Homo* | *sapiens* | ODAM |
| UPI000BADC9C8 | *Ictidomys* | *tridecemlineatus* | ODAM |
| UPI001E26755C | *Lemur* | *catta* | ODAM |
| UPI0003EDE888 | *Leptonychotes* | *weddellii* | ODAM |
| UPI0001C5F341 | *Loxodonta* | *africana* | ODAM |
| UPI000F2F2276 | *Microtus* | *ochrogaster* | ODAM |
| UPI0013438182 | *Moschus* | *moschiferus* | ODAM |
| UPI0000217CB9 | *Mus* | *musculus* | ODAM |
| UPI001C5CA009 | *Orycteropus* | *afer* | ODAM |
| UPI0023FD8446 | *Ovis* | *aries* | ODAM |
| UPI0001F17692 | *Ailuropoda* | *melanoleuca* | SERPINA1 |
| UPI0000124FD5 | *Bos* | *taurus* | SERPINA1 |
| UPI0003C8A3EF | *Camelus* | *ferus* | SERPINA1 |
| UPI0006B11B66 | *Capra* | *hircus* | SERPINA1 |
| UPI0003344B0A | *Condylura* | *cristata* | SERPINA1 |
| UPI000328DF56 | *Dasypus* | *novemcinctus* | SERPINA1 |
| UPI000BB9AFFB | *Delphinapterus* | *leucas* | SERPINA1 |
| UPI0003340CFE | *Echinops* | *telfairi* | SERPINA1 |
| UPI0000F624DC | *Erinaceus* | *europaeus* | SERPINA1 |
| UPI0002990549 | *Felis* | *catus* | SERPINA1 |
| UPI000000CBEC | *Homo* | *sapiens* | SERPINA1 |
| UPI000BADC12A | *Ictidomys* | *tridecemlineatus* | SERPINA1 |
| UPI00123EA0AD | *Leptonychotes* | *weddellii* | SERPINA1 |
| UPI000C81200E | *Loxodonta* | *africana* | SERPINA1 |
| UPI001346A56B | *Moschus* | *moschiferus* | SERPINA1 |
| UPI001C5C8F61 | *Orycteropus* | *afer* | SERPINA1 |
| UPI0000124FDF | *Ovis* | *aries* | SERPINA1 |
| UPI00059B1C44 | *Ailuropoda* | *melanoleuca* | SERPINC1 |
| UPI0000D89868 | *Bos* | *taurus* | SERPINC1 |

| | | | |
|---|---|---|---|
| UPI0003C81BE1 | *Camelus* | *ferus* | SERPINC1 |
| UPI0003AF37D9 | *Capra* | *hircus* | SERPINC1 |
| UPI0003346AD1 | *Condylura* | *cristata* | SERPINC1 |
| UPI000329100E | *Dasypus* | *novemcinctus* | SERPINC1 |
| UPI000BB7B261 | *Delphinapterus* | *leucas* | SERPINC1 |
| UPI000D17FE07 | *Desmodus* | *rotundus* | SERPINC1 |
| UPI001E1DD6E9 | *Echinops* | *telfairi* | SERPINC1 |
| UPI001234CC0A | *Equus* | *caballus* | SERPINC1 |
| UPI0000F61FCD | *Erinaceus* | *europaeus* | SERPINC1 |
| UPI001D19D3E2 | *Felis* | *catus* | SERPINC1 |
| UPI000002C0C1 | *Homo* | *sapiens* | SERPINC1 |
| UPI00025DCD96 | *Ictidomys* | *tridecemlineatus* | SERPINC1 |
| UPI001E26AEEC | *Lemur* | *catta* | SERPINC1 |
| UPI0003EDD592 | *Leptonychotes* | *weddellii* | SERPINC1 |
| UPI000C813313 | *Loxodonta* | *africana* | SERPINC1 |
| UPI00038BE024 | *Microtus* | *ochrogaster* | SERPINC1 |
| UPI001343A7C5 | *Moschus* | *moschiferus* | SERPINC1 |
| UPI0000000900 | *Mus* | *musculus* | SERPINC1 |
| UPI00045E2888 | *Orycteropus* | *afer* | SERPINC1 |
| UPI0000125B63 | *Ovis* | *aries* | SERPINC1 |
| UPI0001F17B27 | *Ailuropoda* | *melanoleuca* | TUFT1 |
| UPI0000136C9D | *Bos* | *taurus* | TUFT1 |
| UPI0003C81F4D | *Camelus* | *ferus* | TUFT1 |
| UPI0003AFE16D | *Capra* | *hircus* | TUFT1 |
| UPI0006428DAE | *Condylura* | *cristata* | TUFT1 |
| UPI00019A5FB8 | *Dasypus* | *novemcinctus* | TUFT1 |
| UPI000BB87305 | *Delphinapterus* | *leucas* | TUFT1 |
| UPI0023810BC5 | *Desmodus* | *rotundus* | TUFT1 |
| UPI00122F45DF | *Echinops* | *telfairi* | TUFT1 |
| UPI0001796107 | *Equus* | *caballus* | TUFT1 |
| UPI0004444238 | *Erinaceus* | *europaeus* | TUFT1 |
| UPI0005ABE932 | *Felis* | *catus* | TUFT1 |
| UPI0000037BFA | *Homo* | *sapiens* | TUFT1 |
| UPI00025DBEA6 | *Ictidomys* | *tridecemlineatus* | TUFT1 |
| UPI001E269DF7 | *Lemur* | *catta* | TUFT1 |
| UPI0003EE1537 | *Leptonychotes* | *weddellii* | TUFT1 |
| UPI0001C5FAD3 | *Loxodonta* | *africana* | TUFT1 |
| UPI00038BB892 | *Microtus* | *ochrogaster* | TUFT1 |
| UPI00134AC1FD | *Moschus* | *moschiferus* | TUFT1 |
| UPI0000027DA2 | *Mus* | *musculus* | TUFT1 |
| UPI001C5CAFE4 | *Orycteropus* | *afer* | TUFT1 |
| UPI00100DA457 | *Ovis* | *aries* | TUFT1 |

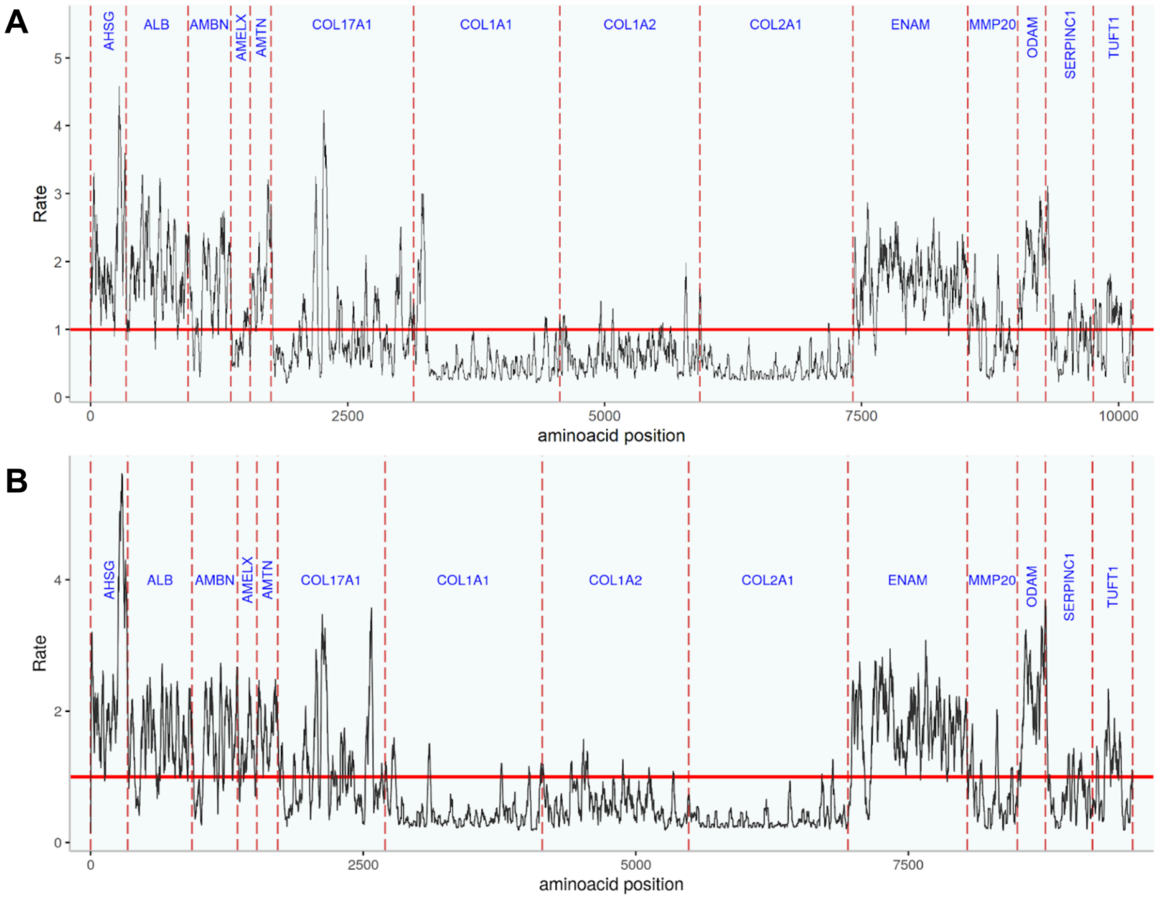

**Supplementary figure S4: Rate4 site scores calculated for a set of 22 mammals**

A - mammals, B - primates (as in Figure 2 in main text)

### 4. Phylogenetic analyses with simulated fragmentary patterns

The predicted protein sequences were fragmented by removing sites (columns of the MSA) to emulate ancient data (see methods in main text for detailed description). From these fragmented datasets, we calculated phylogenetic trees using ML. The topologies of the resulting trees were compared to the reference tree (Kuderna et al. 2023) and RF-distances measured (see results in main text). The differences between the protein based trees and the reference tree were also visually inspected and are described in supplementary table S5. The names of the fragmentation stages roughly represent the age of data from subtropical and tropical fossils they are based on in the model.

**Supplementary table S5** Differences between protein-based trees and the reference tree (Kuderna et al. 2023) at genus level or higher.

| MSA (Protein) | Genus | Reference tree | Protein tree | Difference at family level or higher |
|---|---|---|---|---|
| 5 proteins (full length, except signal peptide) | Galago | sister to Lorisidae | within Lorisidae | yes |
| | Otolemur | sister to Lorisidae | within Lorisidae | yes |
| | Galagoides | sister to Lorisidae | within Lorisidae | yes |
| | Cebuella | sister to Mico + Callibella | sister to Callithrix + Mico + Callibella | no |
| | Semnopithecus | sister to Trachypithecus | unresolved with Trachypithecus | no |
| | Lophocebus | sister to Papio | sister to Mandrillus + Cercocebus | no |
| | Theropithecus | sister to Papio + Lophocebus | sister to Papio + Lophocebus + Mandrillus + Cercocebus | no |
| | Miopithecus | sister to Cercopithecus | sister to all other Cercopithecini | no |
| | Chlorocebus | sister to Allochrocebus | sister to Cercopithecus + Erythrocebus + Allenopithecus + Allochrocebus | no |
| | Allenopithecus | sister to Allochrocebus + Chlorocebus + Erythrocebus | sister to Allochrocebus | no |
| 10 proteins (full length, except signal peptide) | Nasalis | sister to Pygathrix + Rhinopithecus | sister to Pygathrix | no |
| | Semnopithecus | sister to Trachypithecus | unresolved with Trachypithecus | no |
| | Theropithecus | sister to Papio + Lophocebus | sister to Papio | no |
| | Miopithecus | sister to Cercopithecus | sister to all other Cercopithecini | no |
| 14 proteins (full length, except signal peptide) | Carlito | sister to Simiiformes | sister to Strepsirrhini | yes |
| | Cephalopachus | sister to Simiiformes | sister to Strepsirrhini | yes |
| | Tarsius | sister to Simiiformes | sister to Strepsirrhini | yes |
| | Nasalis | sister to Pygathrix + Rhinopithecus | sister to Pygathrix | no |
| | Lophocebus | sister to Papio | sister to Mandrillus | no |
| fragmentation stage 100ka | Carlito | sister to Simiiformes | sister to Strepsirrhini | yes |
| | Cephalopachus | sister to Simiiformes | sister to Strepsirrhini | yes |
| | Tarsius | sister to Simiiformes | sister to Strepsirrhini | yes |
| | Cebuella | sister to Mico + Callibella | sister to Callithrix | no |
| | Lophocebus | sister to Papio | sister to Mandrillus | no |
| | Chlorocebus | sister to Allochrocebus | sister to Erythrocebus | no |
| | Semnopithecus | sister to Trachypithecus | unresolved with Trachypithecus | no |

| | | | | |
|---|---|---|---|---|
| | Rhinopithecus | sister to Pygathrix | sister to Semnopithecus + Trachypithecus | no |
| fragmentation stage 1-2 Ma | Carlito | sister to Simiiformes | sister to all other primates | yes |
| | Cephalopachus | sister to Simiiformes | sister to all other primates | yes |
| | Tarsius | sister to Simiiformes | sister to all other primates | yes |
| | Galago | sister to Lorisidae | within Lorisidae | yes |
| | Otolemur | sister to Lorisidae | within Lorisidae | yes |
| | Galagoides | sister to Lorisidae | within Lorisidae | yes |
| | Aotus | sister to Callitrichidae (weak support in reference (PP = 0.56, all other family nodes PP = 1), thus, discordance not informative) | sister to Atelidae + Cebidae + Callitrichidae | yes |
| | Cebuella | sister to Mico + Callibella | sister to Callithrix | no |
| | Alouatta | together with other Athelidae sister to + Cebidae + Callitrichidae + Aotidae | sister to Callitrichidae | yes |
| | Ateles | together with other Athelidae sister to + Cebidae + Callitrichidae + Aotidae | sister to Callitrichidae | yes |
| | Lagothrix | together with other Athelidae sister to + Cebidae + Callitrichidae + Aotidae | sister to Callitrichidae | yes |
| | Pan | sister to Homo | sister to Gorilla | no |
| | Rhinopithecus | sister to Pygathrix | sister to Semnopithecus + Trachypithecus | no |
| | Chlorocebus | sister to Allochrocebus | sister to Erythrocebus | no |
| | Semnopithecus | sister to Trachypithecus | unresolved with Trachypithecus | no |
| | Mandrillus | sister to Cercocebus | outgroup to Papio + Theropithecus + Lophocebus + Cercocebus | no |
| | Lophocebus | sister to Papio | outgroup to Theropithecus + Papio | no |
| fragmentation stage 5 Ma | Carlito | sister to Simiiformes | sister to Strepsirrhini | yes |
| | Cephalopachus | sister to Simiiformes | sister to Strepsirrhini | yes |
| | Tarsius | sister to Simiiformes | sister to Strepsirrhini | yes |
| | Varecia | basal-most Lemuridae | sister to Lemur + Prolemur + Hapalemur | no |
| | Daubentonia | sister to Indriidae + Lemuridae | sister to Indriidae | yes |
| | Galagoides | sister to Otolemur + Galago | within Galago | no |
| | Arctocebus | sister to Perodictius | within Perodictius | no |
| | all genera of Platyrrhini | NA | Resolution of nodes at family level is widely in discordance with reference tree. | yes |
| | all genera of Catarrhini | NA | Still placed in same family as in reference, but<br>relationships within families with low confidence<br>and mostly in discordance with the reference tree. | no |
| fragmentation stage 10 Ma | all genera of primates | NA | Unresolved phylogeny with bootstrap values typically below 50. Of all infraorders, only Lorisiformes | yes |

| | | | |
|---|---|---|---|
| | | | form a monophylum. |

### 5. Two case studies with simulated fragmentary patterns

Two cases were simulated in order to test how fragmentary protein data can be positioned in a phylogenetic tree, the "Neandertal case" and the "Chimpanzee case".

Three archaic human individuals with sequence data reduced to fragmentary stage "100 ka" were added to a full-length sequence protein MSA of all 14 proteins of interest. This scaffold MSA consisted of five individuals from all other species of *Hominidae* and one hylobatid outgroup. Accordingly, three *Pan troglodyte*s with sequence data reduced to fragmentary stage "5 Ma" were added to the same complete protein scaffold that lacked *Pan troglodyte*s sequences. The modeling of the fragmentation stages is explained in detail in the methods of the main text.

After adding the fragmentary data to the scaffolds, both datasets were aligned with MAFFT v7.520 through the command:

```
mafft --auto --addfragments $fragments --keeplength $scaffold > fragmentary_alignment.fasta
```

ML phylogenetic analysis was performed over the fragmentary alignments with the command:

```
iqtree -nt 4 –s $fragmentary_alignment.fasta -spp $14_prot.nexus -alrt 5000 -bb 5000 –pre $case_out
```

Where the 14_prot_nexus contains the evolutionary models computed for each of the proteins within the concatenation. The resulting trees are displayed in supplementary figure S5.

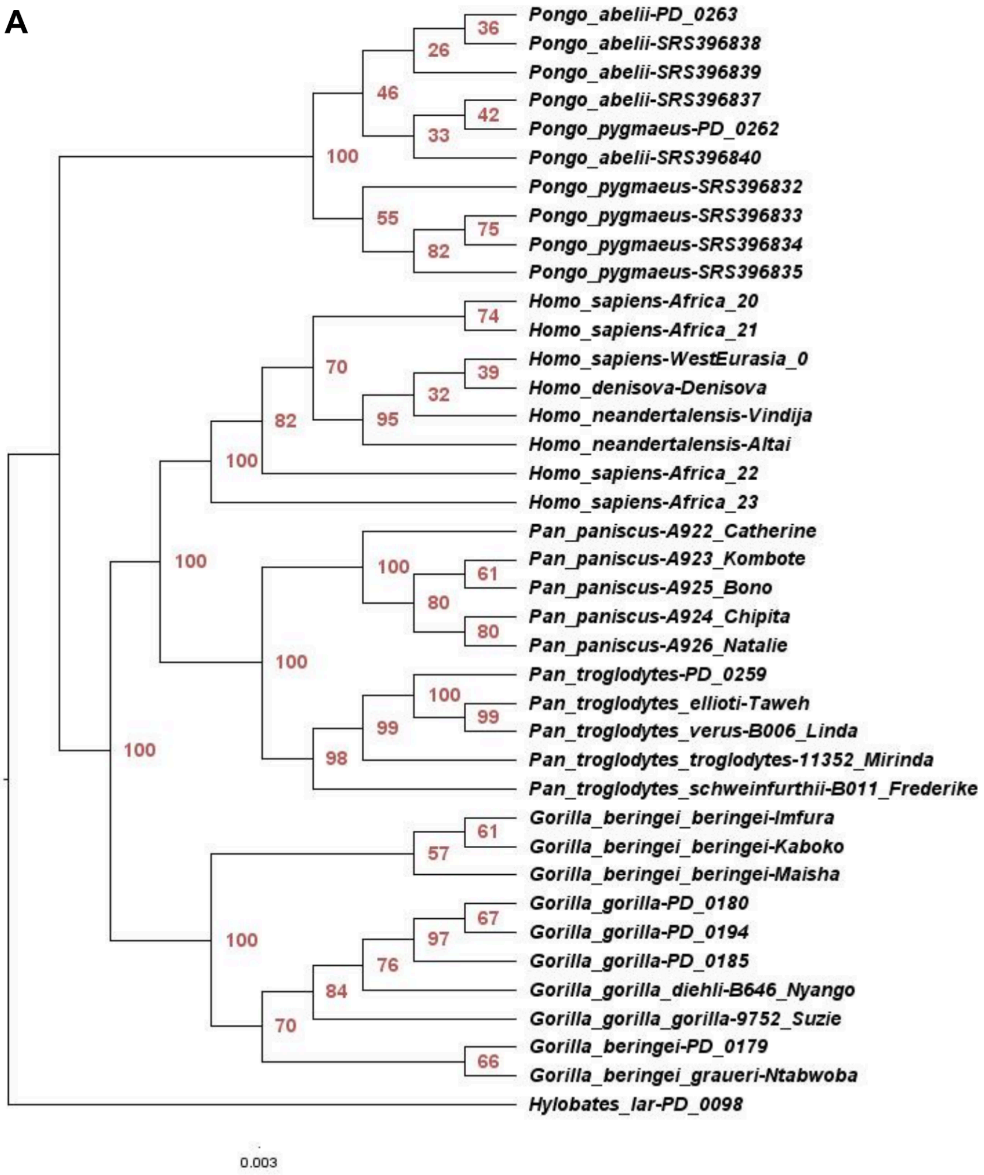

**Supplementary figure S5A: Phylogenetic trees of simulated cases studies**

A “Neanderthal case”; B - “Chimpanzee case”

**B**

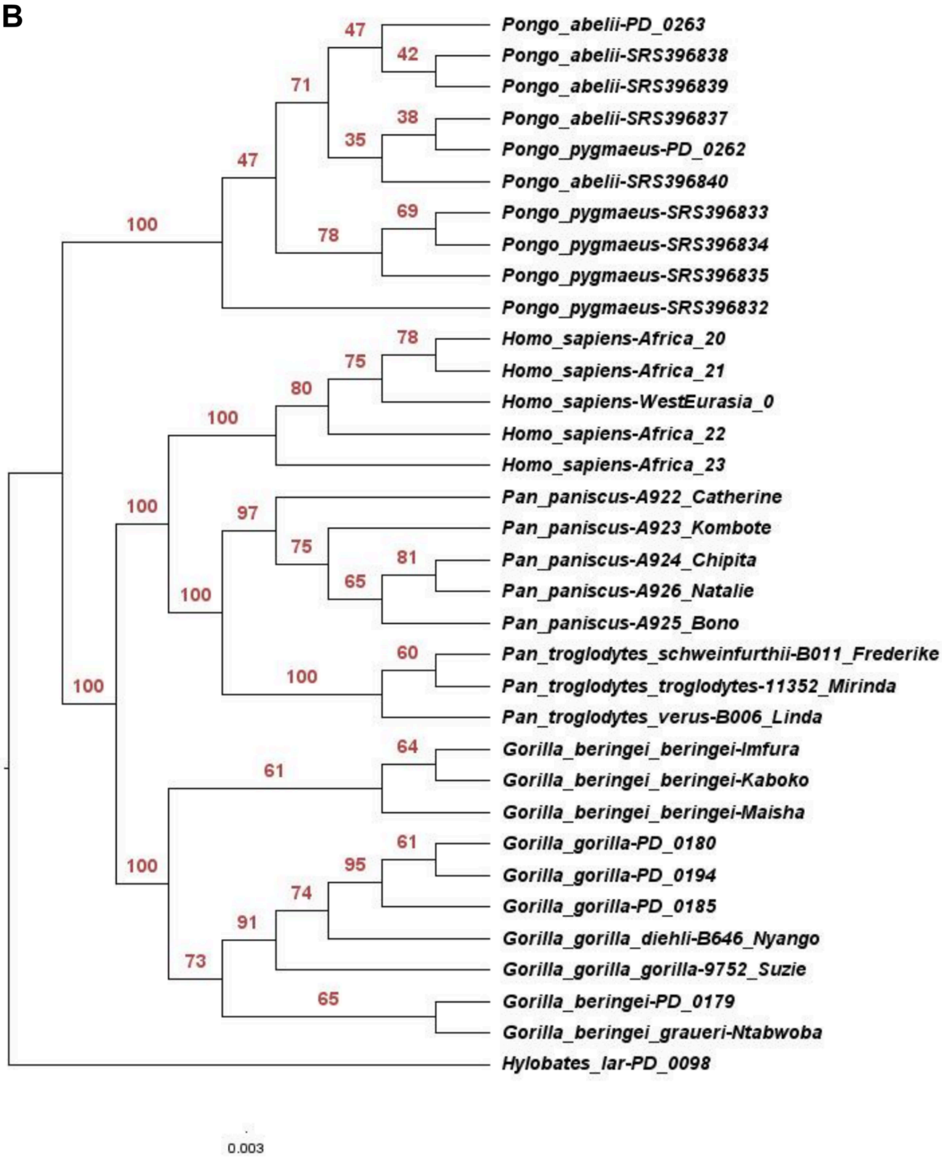

**Supplementary figure S5B: Phylogenetic trees of simulated cases studies**

A “Neanderthal case”; B - “Chimpanzee case”

### 6. Divergence time estimation

The dataset of the “Chimpanzee case” was used to estimate divergence times on a dataset as it could be used in a typical paleoproteomic study. For this, the dataset was reduced to one individual per species and transformed into the sequential phylip format. Divergence times were estimated using fossil calibration with MCMCtree (Yang 2007) of the PAML package v. 4.10.6 (for more details see Methods in main text). Consistency of results was tested in ten independent runs (supplementary figure S6). The estimated divergence times were compared to those in the reference tree (Kuderna et al. 2023) obtained by DNA data (supplementary table S6).

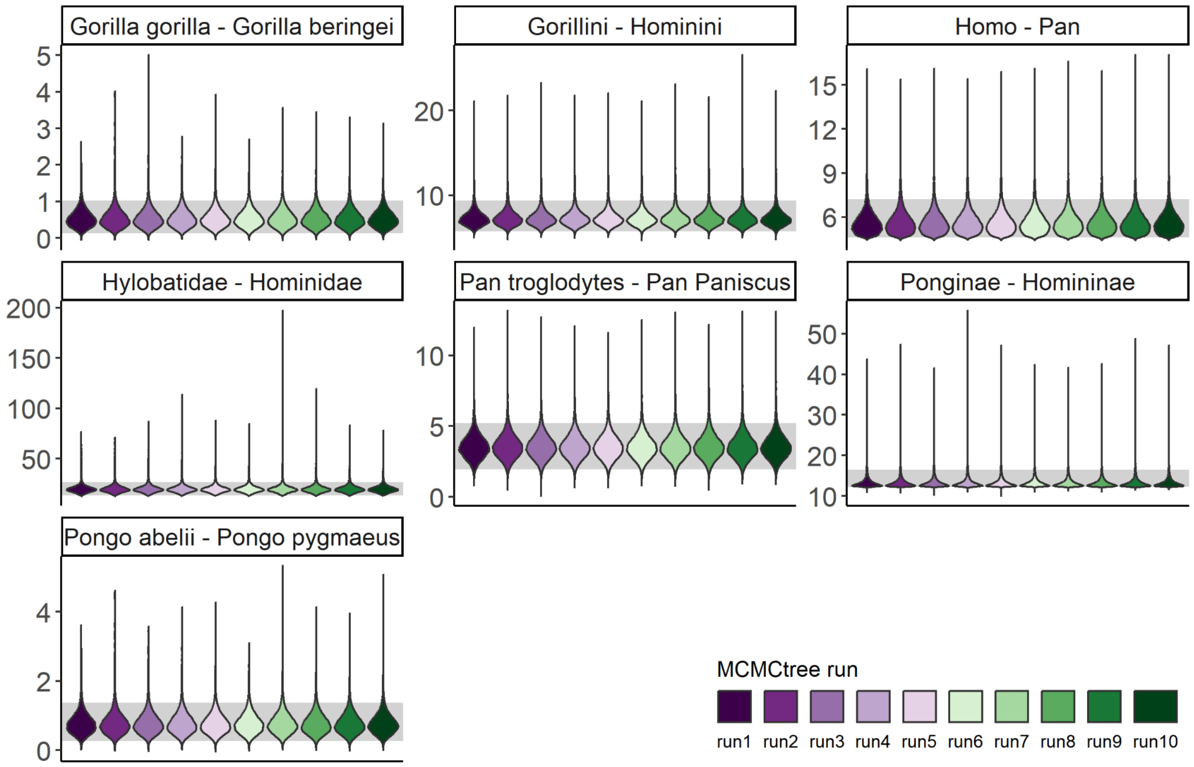

**Supplementary figure S6: Densities of node age estimations**

For each node the posterior densities of node age estimation is shown for 10 independent runs as violin plots. Note the different y-axis scales. The gray area marks the 95% HPD (highest posterior density) for all 10 runs combined.

**Supplementary table S6: Comparison of divergence time estimates at different nodes in Hominidae**

Values for reference from Kuderna et al. (2023). “95% HPD posterior” and “Mean protein” from all 10 Markov chain Monte Carlo runs combined. “Protein” refers to the estimations based on protein sequences from Hominidae (*Pan troglodytes* at fragmentation stage “5 Ma”). “Reference” refers to the estimations based on genomic DNA data from Kuderna et al. (2023).

| **Split** | **Mean protein** | **95% HPD posterior protein** | **95% HPD effective prior protein** | **Mean reference** | **95% HPD posterior reference** | **95% HPD effective prior reference** | **Mean ratio (protein / reference)** |
|---|---|---|---|---|---|---|---|
| Hylobatidae - Hominidae | 20.07 | 13.64 - 26.49 | 13.31 - 30.38 | 22.89 | 21.02 - 24.80 | 13.34 - 21. 30 | 0.88 |
| Ponginae - Homininae | 13.52 | 12.24 - 16.42 | 12.25 - 15.26 | 20.32 | 18.58 - 22.19 | 12.26 - 14.58 | 0.67 |
| Gorillini - Hominini | 7.39 | 5.74 - 9.35 | 6.62 - 14.62 | 10.12 | 8.79 - 11.24 | 7.50 - 12.18 | 0.73 |
| Homo - Pan | 5.72 | 4.63 - 7.21 | 4.63 - 12.87 | 8.01 | 6.91 - 8.96 | 4.63 - 9.41 | 0.71 |
| *Pongo pygmaeus - P. abelii* | 0.78 | 0.26 - 1.37 | 0.03 - 13.22 | 1.55 | 1.18 - 1.95 | 0 - 11.21 | 0.5 |
| *Gorilla gorilla - G. beringei* | 0.55 | 0.13 - 1.02 | 0 - 12.59 | 1.03 | 0.81 - 1.26 | 0 - 9.45 | 0.53 |
| *Pan troglodytes - P. paniscus* | 3.58 | 1.94 - 5.23 | 0 - 10.01 | 2.39 | 1.98 - 2.81 | 0 - 7.05 | 1.5 |

Diversity from 233 Primate Species.” *Science* 380 (6648): 906–13.

Li, Heng. 2011. “A Statistical Framework for SNP Calling, Mutation Discovery, Association Mapping and Population Genetical Parameter Estimation from Sequencing Data.” *Bioinformatics* 27 (21): 2987–93.

Li, Heng, Bob Handsaker, Alec Wysoker, Tim Fennell, Jue Ruan, Nils Homer, Gabor Marth, Goncalo Abecasis, and Richard Durbin. 2009. “The Sequence Alignment/Map Format and SAMtools.” *Bioinformatics* 25 (16): 2078–79.

Nguyen, Lam-Tung, Heiko A. Schmidt, Arndt von Haeseler, and Bui Quang Minh. 2015. “IQ-TREE: A Fast and Effective Stochastic Algorithm for Estimating Maximum-Likelihood Phylogenies.” *Molecular Biology and Evolution* 32 (1): 268–74.

Pupko, Tal, Rachel E. Bell, Itay Mayrose, Fabian Glaser, and Nir Ben-Tal. 2002. “Rate4Site: An Algorithmic Tool for the Identification of Functional Regions in Proteins by Surface Mapping of Evolutionary Determinants within Their Homologues.” *Bioinformatics* 18 (SUPPL. 1): S71–77.

Rambaut, Andrew, Alexei J. Drummond, Dong Xie, Guy Baele, and Marc A. Suchard. 2018. “Posterior Summarization in Bayesian Phylogenetics Using Tracer 1.7.” *Systematic Biology* 67 (5): 901–4.

Ronquist, F., and J. P. Huelsenbeck. 2003. “MRBAYES 3: Bayesian Phylogenetic Inference under Mixed Models.” *Bioinformatics* 19 (12): 1572–74.

Yang, Ziheng. 2007. “PAML 4: Phylogenetic Analysis by Maximum Likelihood.” *Molecular Biology and Evolution* 24 (8): 1586–91.
